# A mouse-adapted Yezo virus model for antiviral testing in immunocompetent mice

**DOI:** 10.64898/2026.03.03.709471

**Authors:** Wenbo Xu, Yuanzhi Wang, Mingming Pan, Qianqian Tan, Fangyu Jin, Liyan Sui, Yinghua Zhao, Nan Liu, Quan Liu

**Affiliations:** Department of Infectious Diseases and Center for Infectious Diseases and Pathogen Biology, Key Laboratory of Organ Regeneration and Transplantation of the Ministry of Education, State Key Laboratory for Diagnosis and Treatment of Severe Zoonotic Infectious Diseases, Key Laboratory for Zoonosis Research of the Ministry of Education, The First Hospital of Jilin University, Changchun, China; Chinese Medicine Guangdong Laboratory, Guangdong Hengqin, China; Department of Pathogenic Biology, School of Medicine, Shihezi University, Shihezi, Xinjiang Uygur Autonomous Region, China

**Keywords:** Orthonairovirus, Yezo virus, Animal model, Antiviral testing, Ribavirin, Viral adaptation

## Abstract

Emerging tick-borne orthonairoviruses pose a growing public health threat, yet few animal models fully recapitulate human disease. Here, we report the development of a mouse-adapted Yezo virus (MA-YEZV) strain that causes lethal infection in immunocompetent mice, mirroring key clinical features of human infection, including thrombocytopenia, leukopenia, and severe liver injury. Serial passaging of YEZV in C57BL/6J mice selected for 31 non-synonymous mutations, enhancing viral replication and pathogenicity. MA-YEZV exhibited broad tissue tropism, with the highest viral loads in the liver, and induced a robust inflammatory response marked by elevated proinflammatory cytokines (e.g., IFN-γ, TNF-α, IL-6) and inflammasome activation. Ribavirin treatment effectively suppressed viral replication, prevented mortality, and mitigated tissue damage, whereas remdesivir showed no efficacy. This model provides a critical tool for studying YEZV pathogenesis and antiviral development, while the identified mutations offer insights into viral adaptation and virulence mechanisms. Our findings underscore the potential of MA-YEZV as a platform for evaluating countermeasures against emerging orthonairoviruses.

## Introduction

The family *Nairoviridae* consists of segmented negative-sense RNA viruses, encompassing at least 51 species across seven genera^1^. These viruses are primarily maintained in arthropods and transmitted to mammals or birds via ticks^2^. Notably, several members of this family are established as significant human and/or animal pathogens, including Crimean-Congo hemorrhagic fever virus (CCHFV), Nairobi sheep disease virus, Dugbe virus, Kasokero virus, Issyk-Kul virus, and Erve virus^3-8^. Recent intensified surveillance has uncovered novel tick-borne orthonairoviruses, such as Tacheng tick virus 1, Songling virus, Yezo virus (YEZV), Wetland virus, and Xue-Cheng virus^9-14^. However, the pathogenic potential of many newly identified viruses remains unverified in laboratory models, hindering the development of targeted therapeutic strategies.

Yezo virus (YEZV), the first human pathogen identified in the Sulina genogroup, was first isolated in 2021 from febrile patients in Hokkaido, Japan^11^. Subsequent cases in Japan and China highlight its emerging public health significance. Clinical manifestations of human infection include fever, headache, dizziness, malaise, lumbago, cough, leukopenia, thrombocytopenia, and elevated liver transaminase activities^15,16^. Additional symptoms may involve gastrointestinal (50%) and neurological (28%) symptoms^16^. *Ixodes persulcatus* ticks are considered the primary vector for YEZV, though the virus has also been detected in *Haemaphysalis megaspinosa* and *Ixodes ovatus* ticks in Japan. Notably, YEZV can disperse via migratory flyways of birds infested with *I. persulcatus*^11,15^. Given the broad distribution of *I. persulcatus* across Asia and Eastern Europe, the geographic range of YEZV likely extends beyond China and Japan^17^.

Although the AG129 mouse model, deficient in type I and II interferon receptors, recapitulates YEZV-induced liver damage in humans, characterized by inflammatory cell infiltration and hepatocyte degeneration, its rapidly progressive lethal disease does not accurately mirror the clinical course in humans^18^. Notably, the lack of interferon signaling in this model impedes the investigation of innate immunity, compromises adaptive immune responses, and restricts analysis of host reactions to infection or vaccination^19^.

In this study, we generated a mouse-adapted virus strain, MA-YEZV, through serial *in vivo* passaging. MA-YEZV induces lethal infection in immunocompetent mice and recapitulates clinical features of human YEZV disease. Critically, ribavirin treatment can protect mice from disease and death. This model provides a valuable platform for investigating YEZV pathogenesis and screening antiviral agents.

## Results

### Generation and characterization of mouse-adapted YEZV (MA-YEZV)

The parental YEZV strain T-HLJ1 establishes only a transient, subclinical infection in immunocompetent C57BL/6J mice, limiting its utility for pathogenicity and transmission studies (Supplementary Fig. 1a-1c)^15^. To overcome this, we generated a mouse-adapted YEZV (MA-YEZV) by serially passaging the tick-derived isolate T-HLJ01 in C57BL/6J mice via intraperitoneal inoculation. Three-week-old female mice were selected based on preliminary findings showing peak hepatic viral loads at 3 days post-infection (dpi; Supplementary Fig. 1d). Serial passages were initiated using clarified liver homogenates from infected mice collected at 3-day intervals (Fig. 1a). Notable signs of adaptation emerged by passage 22 (P22), with mild hepatic jaundice observed at necropsy despite an absence of overt clinical signs. By P30, infected mice developed severe disease manifestations, including piloerection, lethargy, reduced activity, and hypothermia, accompanied by mortality, which prompted a reduction in the passage intervals to 2.5 days. A virus isolate from liver tissue at P40 was designated MA-YEZV for characterization. Hepatic viral titers increased sharply from P0 to P30 and then stabilized between P30 and P40, indicating that the attainment of a stable adaptive state in immunocompetent mice (Fig. 1b). *In vitro* analyses revealed significantly enhanced replication of MA-YEZV compared to T-HLJ01. At multiplicities of infection (MOIs of 0.1 and 0.01), MA-YEZV produced elevated viral RNA levels, higher viral titers, and a greater proportion of antigen-positive cells (Fig. 1c-1e). This enhanced fitness was MOI-dependent, as both strains showed comparable infection efficiency at a high MOI (1.0). To assess potential changes in host tropism, we compared their replication in human cell lines. MA-YEZV demonstrated enhanced replication in human embryonic kidney cells (HEK 293T), lung carcinoma epithelial cells (A549), and hepatocellular carcinoma cells (Hep G2) relative to the parental strain (Supplementary Fig. 2a-2f), suggesting an expanded capacity for human cell infection. Notably, both strains replicated most efficiently in Hep G2 cells, corroborating the pronounced hepatotropism of YEZV observed *in vitro*.

**Fig. 1.**
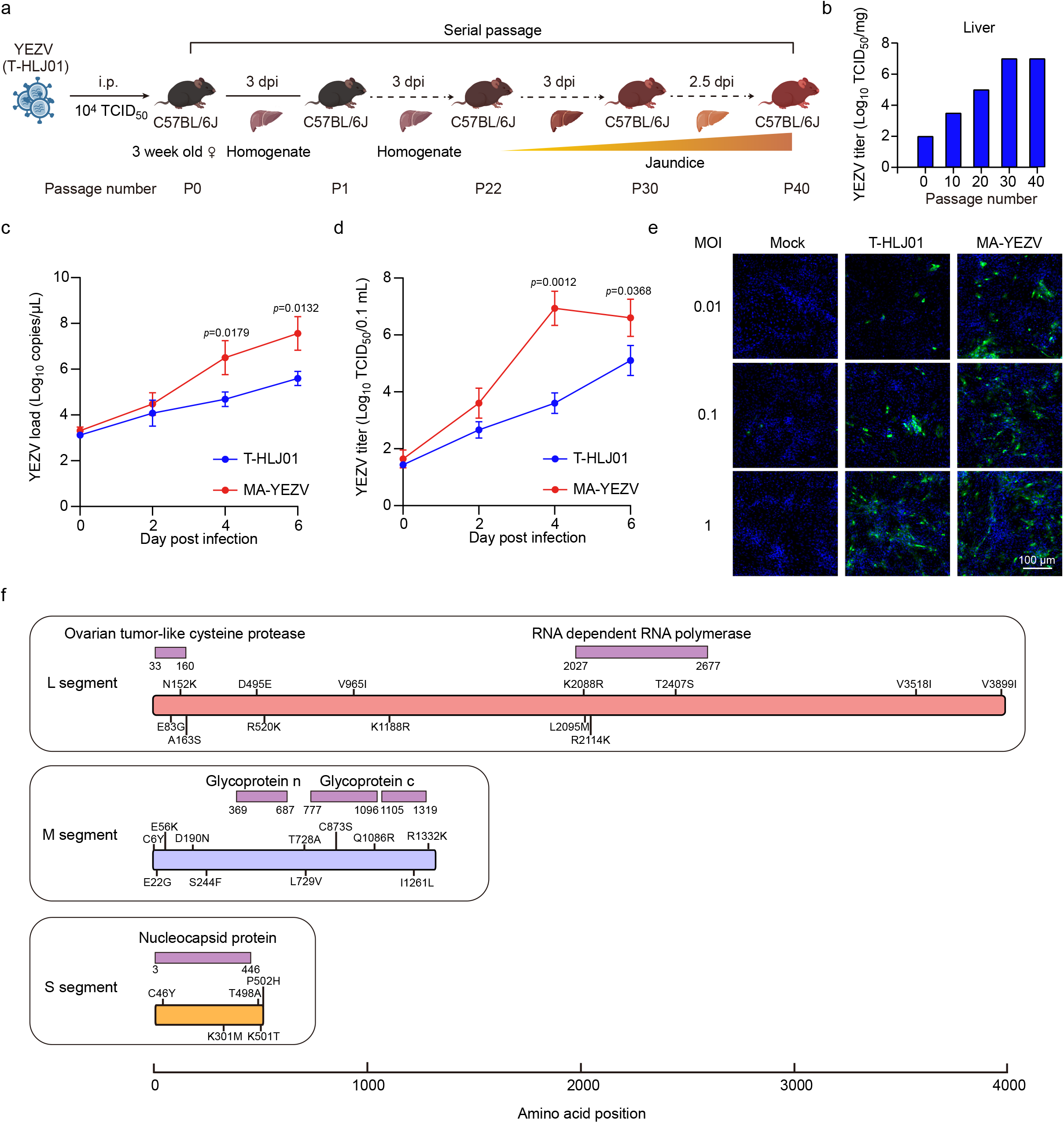
Generation and characterization mouse-adapted YEZV (MA-YEZV) (a) Schematic representation of the viral adaptation process in mice. Three-week-old female C57BL/6J mice were intraperitoneally inoculated with 10^4^ TCID_50_ of the T-HLJ01 strain. Every three days, liver tissues from infected mice were collected, homogenized, centrifuged, and filtered before being passaged into naïve mice. From passage 30, the interval between passages was shortened to 2.5 days. At passage 40, virus was isolated from the liver tissue of infected mice. (b) Viral titers in mouse liver (n=1 independent mice per time point) at passages 0, 10, 20, 30, and 40. (c, d) Comparison of viral RNA copies (c) and infectious titers (d) for T-HLJ01 and MA-YEZV in the supernatants of Vero cells over a 7-day period. Data are presented as mean ± standard deviation (n=3 biological replicates). Statistical significance between two groups at each time point was determined using a two-sided unpaired Student’s t-test. Exact *p* values are indicated in the figure. (e) Immunofluorescence images of Vero cells infected with T-HLJ01 or MA-YEZV at the indicated multiplicity of infection (MOI). Cells were stained for YEZV nucleoprotein (NP) (green) and DAPI (blue) to visualize nuclei. Scale bar, 100 µm. Experiments were performed on three independent biological replicates, and representative images are shown. (f) Analysis of amino acid mutations in the MA-YEZV compared to the parental T-HLJ01 strain. Source data are provided as a Source Data file.

Viral genome analysis revealed that serial passaging led to the accumulation of genetic mutations in YEZV (Fig. 1f, Supplementary Tables 1, 2). The full-length genome sequence of YEZV has been deposited in the GenBank database under accession numbers PV976754-PV976756. Notably, synonymous substitutions accounted for 91.2% of all mutations (312/342). The MA-YEZV genome acquired 31 non-synonymous substitutions, resulting in 13, 11, and 5 amino acid changes in the large (L), medium (M), and small (S) segments, respectively (Supplementary Table 2). Of these mutations, 20 were present in both previously documented human-derived and tick-derived YEZV strains, while 10 were more prevalent in human strains (Supplementary Table 2). Further analysis identified E83G and N152K substitutions in the L segment within the ovarian tumor-like cysteine protease domain (IPR003323). Additional substitutions (K2088R, L2095M, R2114K, and T2407S) located to the Bunyavirus RNA-dependent RNA polymerase (RdRp) domain (IPR007322). No mutations were detected in the Nairovirus Gn (IPR048801) of the M segment. However, three substitutions (C873S, Q1086R, and I1261L) were found in the Gc (IPR002532 and IPR048791). Within the S segment, mutations C46Y and K301M were localized to the nucleocapsid protein (IPR003486). Collectively, these coding-region changes may affect viral assembly and release, receptor-binding affinity, and immune evasion.

Analysis of the untranslated regions (UTRs) identified additional mutations: 10 in the 5′-UTR and 1 in the 3′-UTR of the L segment, 1 in each UTR of the M segment, and 12 in the 3′-UTR of the S segment (Supplementary Table 1). However, these UTR mutations did not substantially alter the predicted RNA secondary structures (data not shown), suggesting that they are unlikely to be the primary drivers of enhanced replication and pathogenicity in MA-YEZV.

### MA-YEZV causes lethal infection across laboratory mouse strains

To evaluate *in vivo* pathogenicity, male and female C57BL/6J mice were intraperitoneally inoculated with 10^4^, 10^2^, and 1 TCID_50_ MA-YEZV, respectively (Fig. 2a). Both the 10^2^ and 10^4^ TCID_50_ doses induced severe disease, with significant weight loss and 100% mortality (12/12) (Fig. 2b, 2c). The median survival time was 7 days for both sexes at 10^4^ TCID_50_, while at 10^2^ TCID_50_, females survived a median of 9 days and males 8 days. Notably, even the 1 TCID_50_ resulted in 91.7% (11/12) mortality in females and 83.3% (10/12) in males, with median survival times of 11.5 and 11 days, respectively, indicating no significant sex-specific differences across doses.

**Fig. 2.**
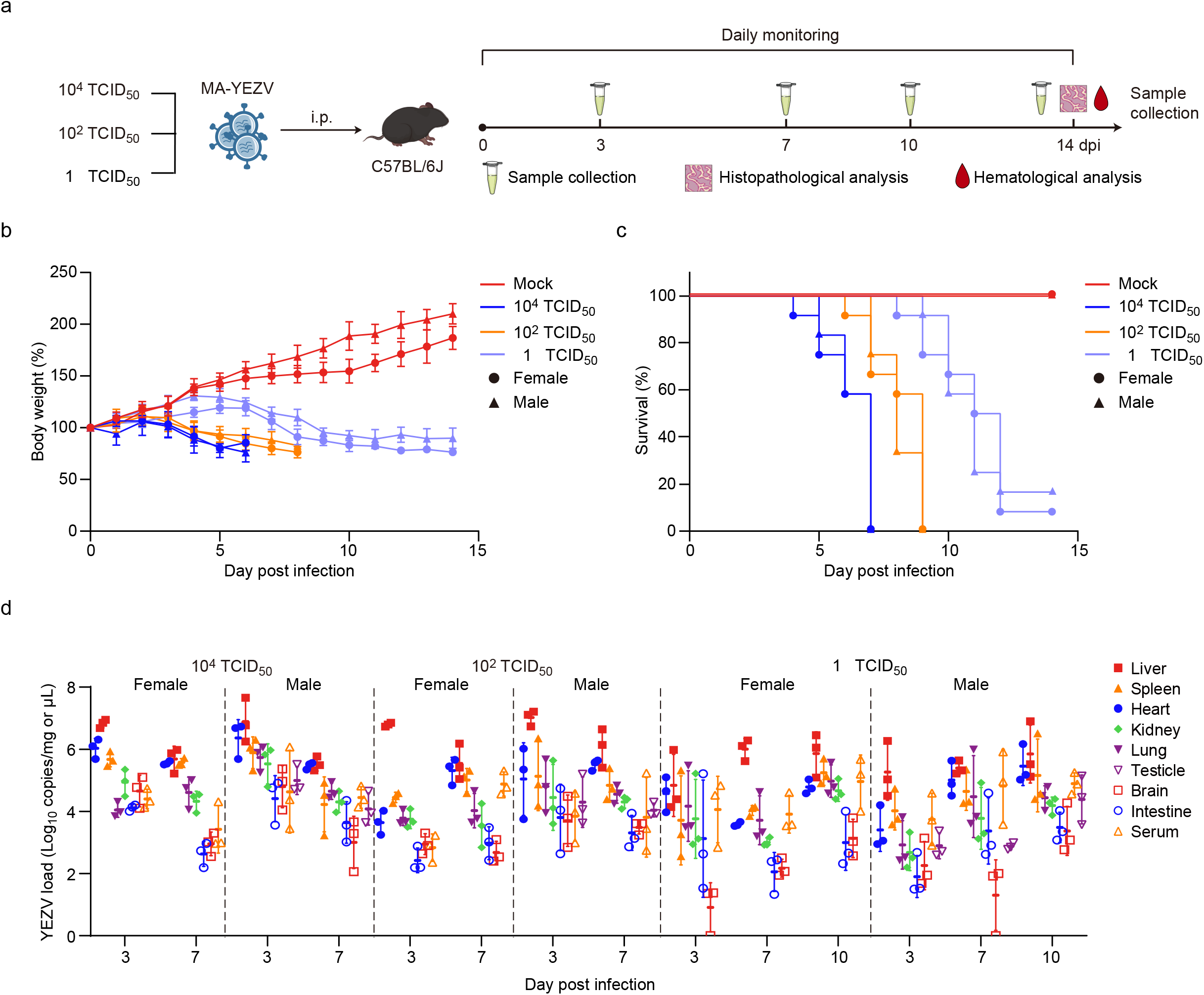
MA-YEZV induces lethal infection in immunocompetent C57BL/6J mice. (a) Schematic representation of MA-YEZV infection in C57BL/6J mice. Three-week-old female and male C57BL/6J mice were intraperitoneally inoculated with MA-YEZV at different doses (10^4^, 10^2^, or 1 TCID_50_). Body weight and survival were monitored daily. At designated time points, tissues and serum were collected for viral load analysis. (b, c) Relative changes in body weight (b) and survival rates (c) of MA-YEZV-infected mice (n=12 independent mice per group) compared to mock-infected controls (n=6 independent mice). (d) Viral loads in tissues and serum of intraperitoneally infected mice (n=3 biological replicates) measured by RT-qPCR at 3, 7, and 10 days post-infection. Data are presented as mean ± standard deviation (b, d). Source data are provided as a Source Data file.

RT-qPCR showed that serum viral loads remained consistent across dose groups during infection, ranging from ∼10^4.12^ copies/mg at 3 dpi to ∼10^4.04^ copies/mg at 10 dpi. The liver consistently exhibited the highest viral loads among tissues. In the high- and medium-dose groups, hepatic viral loads peaked early (10^6.83^ copies/mg for high dose; 10^6.89^ copies/mg for medium dose), but declined significantly by 7 dpi (10^5.70^ copies/mg for high dose; 10^5.80^ copies/mg for medium dose). In contrast, the low-dose group showed a sustained increase in hepatic viral load, rising from ∼10^5.05^ copies/mg at 3 dpi to ∼10^5.85^ copies/mg (Fig. 2d). Paradoxically, despite increasing viral loads in the 1 TCID_50_ group, viral titers decreased after 7 dpi (Supplementary Fig. 3), consistent with observations in a CCHFV-infected mouse model^20^.

To determine whether viral adaptation altered tissue tropism, we compared viral titers in tissues and serum at 3 dpi between mice infected with the parental T-HLJ01 strain and those infected with MA-YEZV. In both groups, viral titers were highest in the liver, indicating that core hepatic tropism was preserved following serial passage (Supplementary Fig. 3). However, MA-YEZV infection resulted in significantly elevated viral titers in other tissues and in serum compared with T-HLJ01 (Supplementary Fig. 3). These findings suggest while the primary tissue tropism remained unchanged, mouse adaptation markedly enhanced systemic infectivity of MA-YEZV. Viral RNA levels in other tissues were uniformly low, though infectious virus could still be isolated from the serum, spleen, and testes at 7 dpi. Given the 1 TCID_50_ dose did not cause universal mortality and showed no sex bias, it was selected for subsequent infections in male mice.

To evaluate route dependency, male C57BL/6J mice were inoculated with 1 TCID_50_ of MA-YEZV subcutaneously or intranasally (Supplementary Fig. 4a). Both routes caused lethal infection, but mortality rates were lower than with those for intraperitoneal injection: 75.0% (9/12) with subcutaneous (s.c.) inoculation (median survival, 11 days) and 66.7% (8/12) with intranasal (i.n.) inoculation (median survival, 13 days) (Supplementary Fig. 4b, 4c). RT-qPCR showed the liver had the highest viral RNA levels regardless of route (10^6.28^ copies/mg for i.n.; 10^5.84^ copies/mg for s.c. at 14 dpi). Subcutaneously inoculated mice exhibited high viral RNA levels in the heart (10^5.47^ copies/mg at 14 dpi) and spleen (10^4.88^ copies/mg at 14 dpi), while intranasally infected mice showed elevated levels in the lungs (10^4.56^ copies/mg at 14 dpi), comparable to those in the heart (10^4.98^ copies/mg at 14 dpi) and spleen (10^4.40^ copies/mg at 14 dpi) (Supplementary Fig. 4d, 4e).

To characterize shedding kinetics, viral RNA was detected early in oropharyngeal and rectal swabs but declined sharply after 6 dpi (Supplementary Fig. 5a). Infectious virus was isolated from oropharyngeal (6/6) and rectal swabs (5/6) at 3 dpi, but by 6 dpi, only 1 out of 6 mice yielded virus from oropharyngeal swabs, and none from rectal swabs (data not shown). For direct contact transmission, male C57BL/6J mice inoculated with 1 TCID_50_ of MA-YEZV (index mice; n=3) were housed individually for 3 days before co-housing with naïve male contacts (n=3) (Supplementary Fig. 5b). In index mice, infectious viral titers in the liver and serum increased during early infection (liver: 10^2.33^ TCID_50_/mg at 3 dpi to 10^4.33^ TCID_50_/mg at 7 dpi; serum: 10^2.33^ TCID_50_/μL at 3 dpi to 10^3.53^ TCID_50_/μL at 7 dpi) before declined markedly at 10 dpi (liver: 10^2.17^ TCID_50_/mg; serum: 10^2.33^ TCID_50_/μL) (Supplementary Fig. 5c). Infectious virus was also detected in oropharyngeal and rectal swabs (∼10^1.33^ TCID_50_/μL) at 3 and 5 dpi, but fell below the detection limit by 7 dpi (Supplementary Fig. 5d). Although index mice harbored and shed infectious virus at levels theoretically sufficient for transmission, no transmission to contact mice was observed. Notably, no infections or viral replication occurred in the contact mice, even after they cannibalized an index mouse that died at 12 dpi, indicating failed transmission despite extreme exposure (Supplementary Fig. 5e, 5f). Additionally, infection of pregnant C57BL/6J mice at embryonic day 12.5 (E 12.5) with MA-YEZV resulted in no detectable YEZV RNA in the offspring (data not shown). These findings indicate that MA-YEZV may lack both contact and vertical transmission capability in immunocompetent mice.

To investigate whether mouse adaptation conferred cross-strain pathogenicity, male BALB/c and Kunming (KM) mice were inoculated with 1 TCID_50_ of MA-YEZV (Supplementary Fig. 6a). Both strains exhibited weight loss and mortality comparable to C57BL/6J mice (Supplementary Fig. 6b, 6c). BALB/c mice were more susceptible, with 100% mortality (12/12) and a median survival of 6 days, while KM mice showed 58.3% mortality (7/12) and a median survival of 10 days. Viremia persisted throughout infection in both mouse strains, peaking at endpoint (10^4.70^ copies/mg in BALB/c mice; 10^4.99^ copies/mg in KM mice). MA-YEZV showed liver tropism in BALB/c and KM mice, similar to C57BL/6J mice: hepatic viral loads increased from 10^4.32^ copies/mg at 3 dpi to 10^5.80^ copies/mg at 7 dpi in BALB/c mice, and more slowly in KM mice from 10^3.90^ copies/mg at 3 dpi to 10^5.64^ copies/mg at 14 dpi. The spleen had the second-highest viral RNA load, with 10^4.76^ copies/mg in BALB/c at 7 dpi and 10^4.68^ copies/mg in KM mice at 14 dpi. Other tissues showed uniformly low viral RNA levels, which increased gradually over infection (Supplementary Fig. 6d). These results establish MA-YEZV as a robust model for lethal infection across common laboratory mouse strains.

### Pathological and biochemical changes induced by the MA-YEZV in mice

To further characterize MA-YEZV-associated pathology, C57BL/6J mice were intraperitoneally inoculated with a low dose (1 TCID_50_) of MA-YEZV to prevent premature mortality. Mice inoculated with a high dose (10^4^ TCID_50_) of the parental T-HLJ01 strain via the same route served as controls. MA-YEZV–infected mice displayed marked hepatic jaundice at 10 dpi (Fig. 3a). Immunohistochemistry revealed progressively increasing viral antigen as infection progressed. Histologically, liver injury was marked by inflammatory cell infiltration, starting as focal lesions and later developing into multifocal hepatocellular necrosis. In contrast, T-HLJ01–infected livers showed only low levels of viral antigen at 3 dpi, with irregular hepatocellular degeneration appearing as single-cell or cord-like lesions that subsequently resolved. Viral antigen was undetectable at later time points, and the mild diffuse hepatocellular degeneration observed at 10 dpi likely reflects inter-individual variability during recovery (Fig. 3a).

**Fig. 3.**
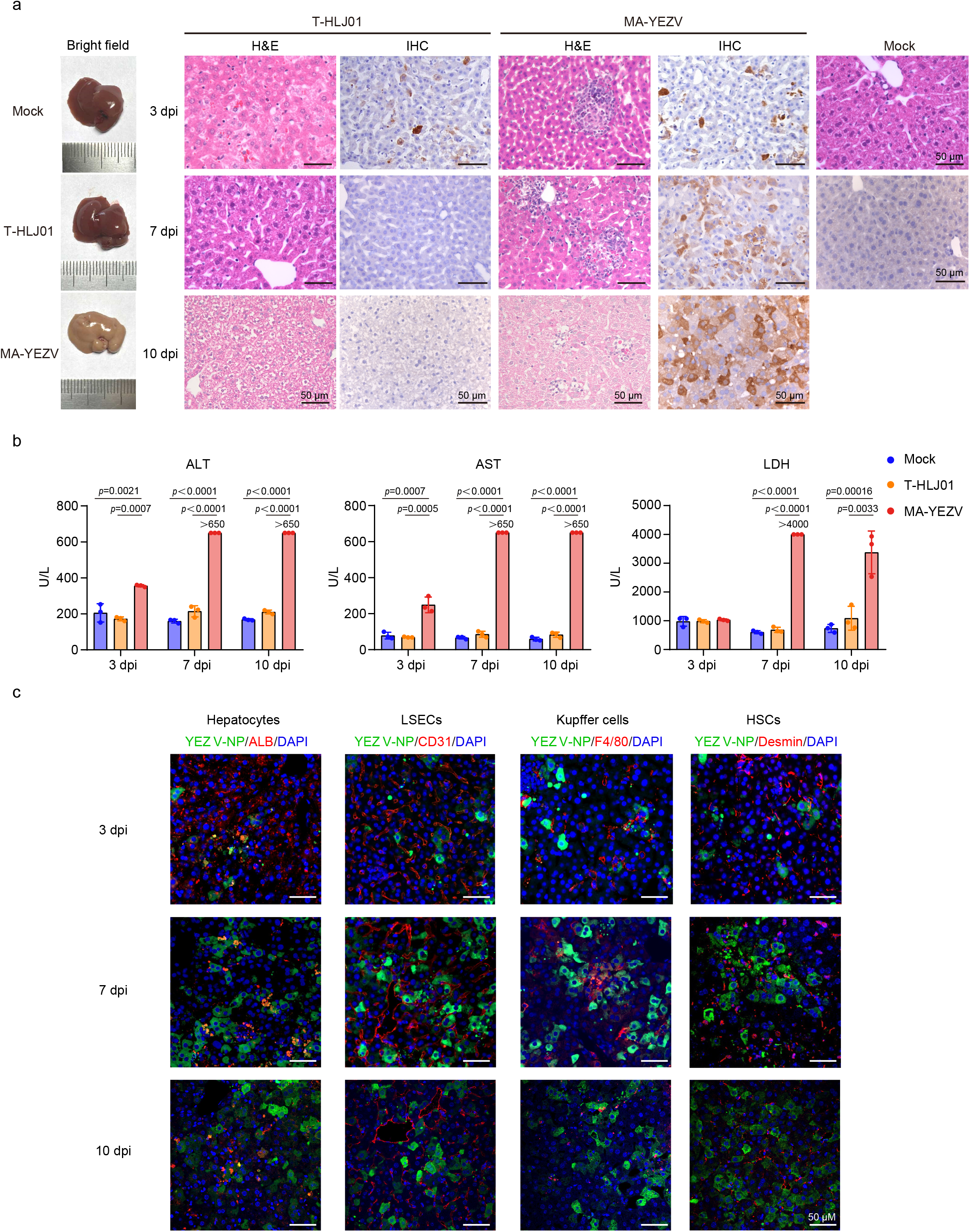
Liver pathology in C57BL/6J mice infected with MA-YEZV. Three-week-old male C57BL/6J mice were intraperitoneally inoculated with 1 TCID_50_ of MA-YEZV or 10^4^ TCID_50_ of the T-HLJ01 strain. Ten days post-infection, liver and blood samples were collected for pathological and biochemical analyses. (a) Liver histopathological assessment, including gross morphology, hematoxylin and eosin (H&E) staining, and immunohistochemistry (IHC) using an antibody against the YEZV nucleoprotein (NP), was performed to evaluate the pathogenicity of MA-YEZV. Scale bars, 100 μm (H&E) and 50 μm (IHC). (b) Serum biochemical markers are indicative of liver injury. Data are presented as mean ± standard deviation (n=3 biological replicates). ALT, alanine aminotransferase; AST, aspartate aminotransferase; LDH, lactate dehydrogenase. Statistical significance was determined by one-way ANOVA (two-sided) followed by Tukey’s multiple-comparison test to adjust for multiple testing. Exact *p* values are provided in the figure legends. (c) Immunofluorescence showing co-localization of YEZV NP (green) with cellular markers: hepatocytes (albumin, ALB; red), liver sinusoidal endothelial cells (LSECs; CD31; red), hepatic stellate cells (HSCs; Desmin; red) and Kupffer cells (F4/80; red). Scale bar, 50 μm. IHC, IF, H&E staining, and bright-field imaging were performed on three independent biological replicates. The images shown are representative of these experiments. Source data are provided as a Source Data file.

Consistent with the histopathological changes, serum biochemistry revealed significant liver injury in MA-YEZV-infected mice. Alanine aminotransferase (ALT) and aspartate aminotransferase (AST) were elevated by 3 dpi, while lactate dehydrogenase (LDH) increased markedly from 7 dpi onward, a pattern mirroring clinical YEZV infection (Fig. 3b).^16^ MA-YEZV infection also reduced serum albumin (ALB), globulin (GLB), and total protein (TP), indicating impaired hepatic synthesis. Furthermore, elevated alkaline phosphatase (ALP), total bile acid (TBA), and uric acid, alongside decreased triglycerides (TG), glucose (GLU), and inorganic phosphorus (IP), reflected broader metabolic dysfunction. T-HLJ01 infection, however, did not significantly alter ALB, GLB, or TP levels and induced only mild, transient changes in ALP and TG that resolved by 10 dpi (Supplementary Fig. 7, Supplementary Table 3, 5, 7).

Hematological analysis showed that MA-YEZV infection recapitulated key clinical features of human disease.^16^ White blood cell and lymphocyte counts were persistently suppressed, and platelet counts dropped significantly from 7 dpi. In contrast, T-HLJ01 infection caused only transient leukopenia and lymphopenia at 3–7 dpi, with a mild platelet reduction observed only at 10 dpi (Supplementary Fig. 7, Supplementary Table 4, 6, 8).

To identify cellular targets, liver sections were co-stained for YEZV nucleoprotein (NP) and specific cellular markers. At 3 dpi, NP colocalized with ALB^+^ hepatocytes and was detected at the periphery and in the cytoplasma of CD31^+^ liver sinusoidal endothelial cells and F4/80^+^ Kupffer cells, but not in the desmin^+^ hepatic stellate cells. By 7 dpi, NP expression was widespread, showing extensive colocalization with hepatocytes and clear detection in all four cell types, including desmin^+^ hepatic stellate cells. At 10 dpi, despite substantial tissue damage, NP remained detectable in hepatocytes, endothelial cells, stellate cells, and the remaining Kupffer cell population (Fig. 3c). These findings demonstrate that YEZV exhibits broad tropism for multiple hepatic cell types.

Beyond the hepatic pathology, systematic evaluation of other tissues (spleen, testicle, kidney, heart, lung, intestine, colon, and brain) revealed additional alterations (Supplementary Fig. 8). MA-YEZV infection was associated with lymphocyte depletion in splenic white pulp, scattered neutrophil infiltration in the red pulp, and subset-specific apoptosis of testicular spermatogenic cells (subset-specific). It also induced goblet cell swelling in the colon and edema in the small intestinal lamina propria (Supplementary Fig. 8). In contrast, T-HLJ01 infection showed no detectable antigen in the spleen or testes and no significant testicular pathology; notably, it induced splenic macrophage proliferation with phagocytosis of apoptotic cells, a tissue repair response that may restrict its replication in immunocompetent mice. The colon and small intestine exhibited no pathological changes following T-HLJ01 infection. Importantly, neither viral strain induced significant pathological changes in the heart, lung, kidney, or brain (Supplementary Fig. 8).

### MA-YEZV induces a robust inflammatory immune response

Nairovirus infections typically elicit elevated levels of inflammatory cytokines^20,21^. To investigate the host inflammatory response, we first assessed viral replication in serum and liver. Serum viral loads did not differ significantly between T-HLJ01 and MA-YEZV-infected mice at 3 dpi. However, by 7 dpi, viral RNA in T-HLJ01-infected mice fell below the detection limit, where the MA-YEZV loads increased significantly increased. A similar divergence was observed in the liver, where MA-YEZV titers were already notably higher than those of T-HLJ01by 3 dpi (Fig. 4a). As nairovirus infections typically induce robust cytokine release, we next analyzed serum inflammatory mediators. Serum cytokine analysis in mice revealed that MA-YEZV infection significantly increased IL-6, IL-10, IFN-γ, MCP-1 (CCL2), and TNF-α at both 3 and 7 dpi, while IL-12p70 showed a non-significant upward trend. In contrast, the parental T-HLJ01 strain did not induce significant increases in these cytokines (Fig. 4b).

**Fig. 4.**
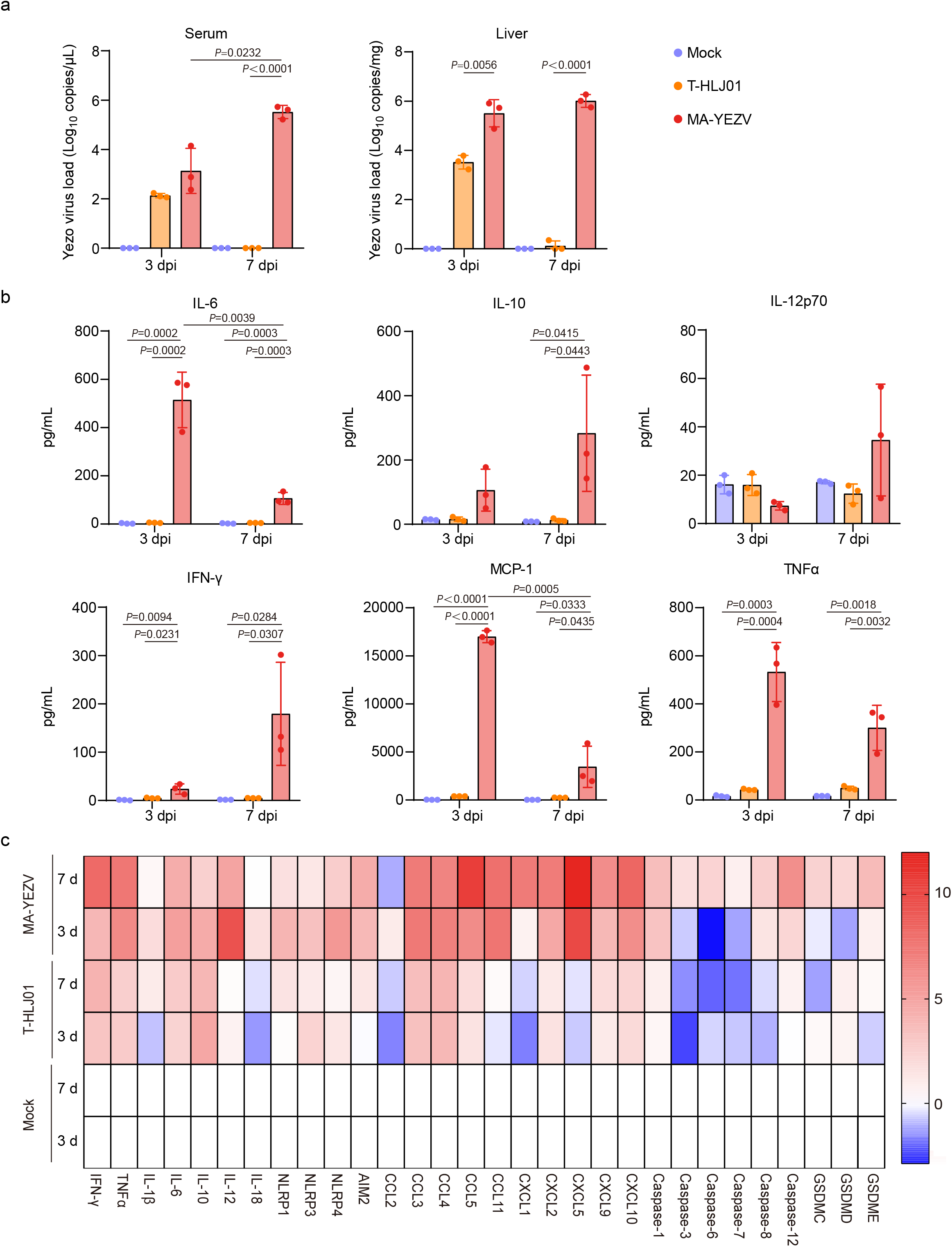
MA-YEZV induces robust inflammatory responses. Three-week-old male C57BL/6J mice were intraperitoneally inoculated with 1 TCID_50_ of MA-YEZV or 10^4^ TCID_50_ of the T-HLJ01 strain. Liver and blood samples were collected at designated time points for cytokine analysis. (a) Viral loads in the liver and serum of mice infected with MA-YEZV or T-HLJ01 strains (n=3, biological replicates) measured by RT-qPCR at 3 and 7 days post-infection (dpi). (b) Serum cytokine profiles assessed by CBA in mice infected with T-HLJ01 or MA-YEZV (n=3, biological replicates) at 3 and 7 dpi. Analytes measured include IL-6, IL-10, IL-12p70, IFN-γ, MCP-1, TNF-α. (c) Hepatic cytokine mRNA expression levels measured by RT-qPCR in MA-YEZV-infected mice (n=3 biological replicates, relative expression). Data are presented as mean ± standard deviation (a, b). Statistical significance between two groups at each time point was determined using a two-sided unpaired Student’s t-test. For comparisons among multiple groups, one-way ANOVA (two-sided) was performed, followed by Tukey’s multiple-comparison test to adjust for multiple testing. Exact *p* values are indicated in the figure. Source data are provided as a Source Data file.

Given the hepatic impairment observed in YEZV-infected mice, we analyzed hepatic cytokine responses (Fig. 4c). Transcript analysis showed that T-HLJ01 induced only modest increases in *IFN-γ, TNF-α, IL-6, IL-10, IL-18, CCL3*, and *CCL4*. In stark contrast, MA-YEZV infection triggered widespread and significant upregulation of multiple cytokines and inflammasome components, including pro-inflammatory cytokines (I*FN-γ, TNF-α, IL-6, IL-12*), immunoregulatory cytokines (*IL-10*), inflammasome sensors (*NLRP1, NLRP3, NLRP4, AIM2*) and chemokines (*CCL3, CCL4, CCL5, CCL11, CXCL1, CXCL2, CXCL5, CXCL9, CXCL10*). Additionally, markers of pyroptosis, including caspase and GSDM family members, were elevated at 7 dpi (Fig. 4c).

Immunoblot analysis revealed distinct temporal patterns of viral nucleoprotein (NP) expression. In T-HLJ01-infected mice, hepatic NP levels peaked at 3 dpi before declining rapidly, whereas in MA-YEZV-infected mice, NP expression remained persistently elevated beyond this time point (Supplementary Fig. 9a). Concordantly, the inflammasome sensor NLRP3 was significantly upregulated in MA-YEZV-infected livers, while only a modest increase was observed following T-HLJ01 infection. These findings indicate that MA-YEZV infection induces a robust inflammatory response and pyroptosis, which may contribute to its associated mortality.

### Ribavirin confers complete protection against lethal MA-YEZV infection

To identify compounds with potential antiviral activity against MA-YEZV, the antiviral effects of ribavirin and remdesivir were evaluated *in vitro* using Vero cells. Ribavirin potently inhibited MA-YEZV replication (IC_50_ = 18.2 μM), whereas neither GS-441524 nor GS-5734 showed any antiviral activity at concentrations up to 200 μM (Supplementary Fig. 10a).

To validate the utility of the mouse challenge model, we assessed the protective efficacy of two broad-spectrum antivirals, ribavirin (HY-B0434) and remdesivir (GS-5734). C57BL/6J mice received intraperitoneally injections of ribavirin (50 or 100 mg/kg) or remdesivir (10 or 30 mg/kg) beginning 2 hours post-infection, with daily dosing for 7 consecutive days (Fig. 5a). Over a 14-day monitoring period, vehicle-treated mice developed significant weight loss, with a mortality rate of 91.7% (11/12) and a median survival time of 12 days. In contrast, ribavirin treatment at both doses completely prevented significant weight loss and conferred 100% survival. Remdesivir, by contrast, lacked protective activity against the mouse-adapted YEZV strain: administration of 10 mg/kg remdesivir resulted in 100% mortality, with a median survival time (11.5 days) marginally shorter than that of the vehicle-treated controls (Fig. 5b, 5c). RT-qPCR analysis revealed persistent MA-YEZV replication in multiple tissues and serum of mice in the vehicle-treated mice, with the highest increase observed in the liver (10^6.57^ copies/mg). Ribavirin treatment potently reduced viral loads, with only trace amounts of viral RNA detected at 3 dpi, and negative results at 7 and 10 dpi. In contrast, remdesivir had negligible efficacy, as viral loads in treated mice were comparable to those in vehicle controls, with sustained replication and the highest loads in the liver (Fig. 5d).

**Fig. 5.**
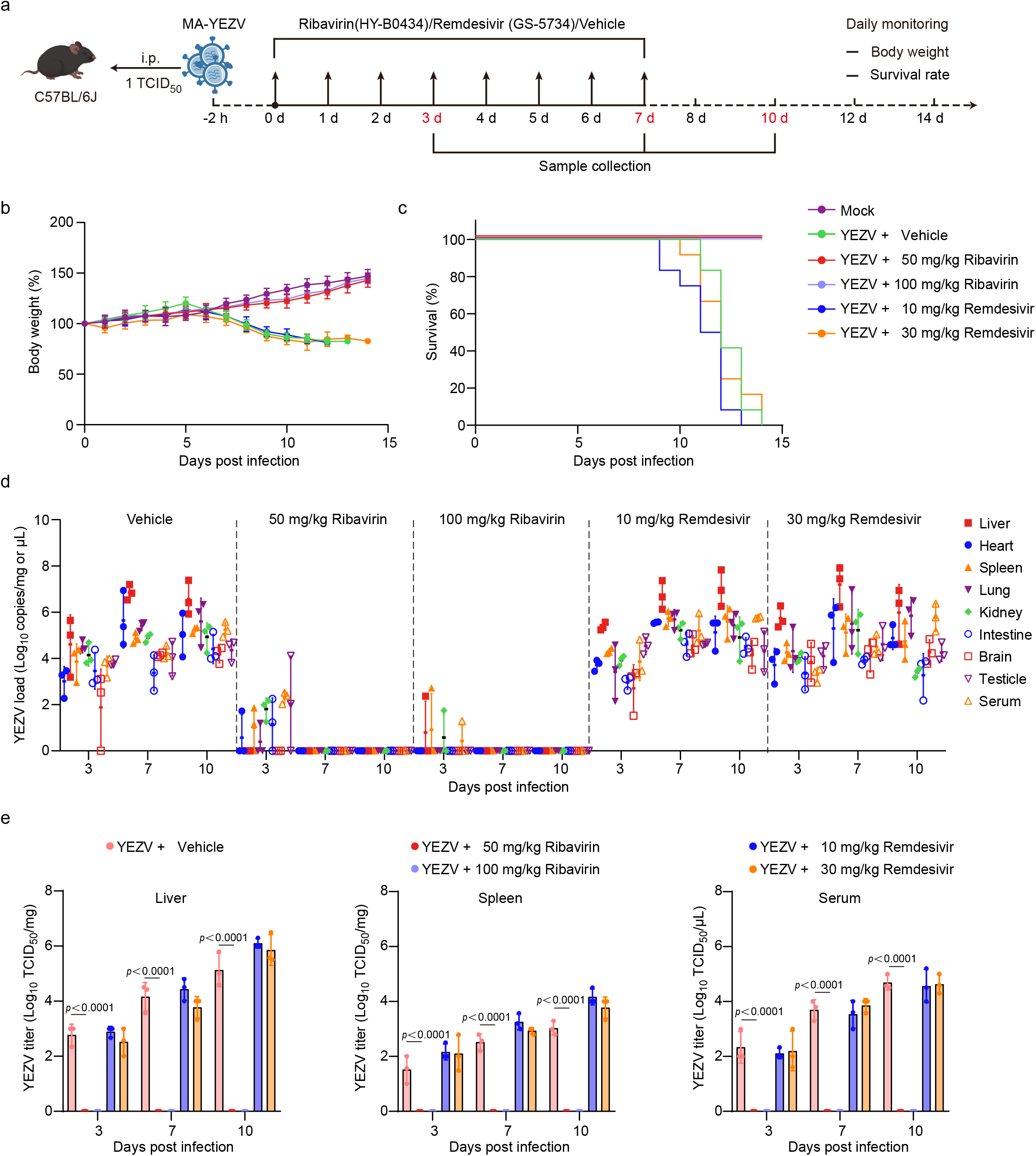
Antiviral efficacy of candidate drugs in MA-YEZV infected C57BL/6J mice. (a) Schematic of the antiviral treatment protocol. Three-week-old male C57BL/6J mice were intraperitoneally inoculated with 1 TCID_50_ of MA-YEZV. Two hours post-infection, mice were treated with either ribavirin (50 or 100 mg/kg) or remdesivir (10 or 30 mg/kg), while vehicle-treated mice served as controls. Antiviral treatments were administered once daily for a total of 8 consecutive days. Body weight and survival were monitored daily. At designated time points, tissues and serum were collected for viral analysis. (b, c) Effects of antiviral treatment on body weight changes (B) and survival rates (C) of MA-YEZV-infected mice (n=12 independent mice per group), compared to vehicle-treated controls (n=6 independent mice). (d) Viral load reduction in tissues and serum of infected mice (n=3 biological replicates) treated with antiviral drugs or vehicle after MA-YEZV infection. (e) Infectious virus titers in the liver, spleen, and serum of treated of infected mice (n=3 biological replicates) treated with antiviral drugs or vehicle after MA-YEZV infection. Data are presented as mean ± standard deviation (b, d, e). Statistical significance was determined by one-way ANOVA (two-sided) followed by Tukey’s multiple-comparison test to adjust for multiple testing. Exact *p* values are provided in the figure legends. Source data are provided as a Source Data file.

Infectious virus titers mirrored viral RNA levels, with peak liver titers in the vehicle-treated mice reaching 10^5.13^ TCID_50_/mg at 10 dpi. Lower but sustained titers were observed in serum (10^4.75^ TCID_50_/mL) and spleen (10^3.03^ TCID_50_/mg) at this time point. Ribavirin treatment eliminated detectable infectious virus in the liver, spleen, and serum by 3, 7, and 10 dpi, whereas remdesivir failed to suppress viral replication, with titers remaining comparable to vehicle controls (Fig. 5e).

The stability and efficacy of remdesivir (GS-5734) in mice are limited by high serum carboxylesterase 1c (Ces1c) activity^22^. We therefore also evaluated its major circulating metabolite, GS-441524, which exhibits more favorable pharmacokinetics in rodents^23^. *In vivo*, GS-441524 proved similarly ineffective as the parent compound, failing to improve survival, reduce viral loads, or mitigate histopathological damage in the liver (Supplementary Fig. 10b-10f). Together, these data demonstrate the potent and specific antiviral efficacy of ribavirin against MA-YEZV, in stark contrast to the inactivity of both remdesivir and its primary metabolite.

### Ribavirin alleviates pathological damage and inflammatory responses in MA-YEZV-infected mice

Pathological analysis further validated ribavirin’s antiviral efficacy. Macroscopic examination revealed significantly reduced hepatic jaundice in ribavirin-treated mice (Fig. 6a). Histologically, livers from vehicle-treated controls exhibited widespread moderate-to-severe hepatocellular swelling, vacuolar degeneration, dilated central/portal veins, enlarged sinusoids, and scattered cellular debris with pyknotic nuclei, accompanied by intense viral antigen staining. In contrast, ribavirin-treated livers closely resembled those of mock-infected mice, with only minimal hepatocellular necrosis and no detectable viral antigens (Fig. 6a).

**Fig. 6.**
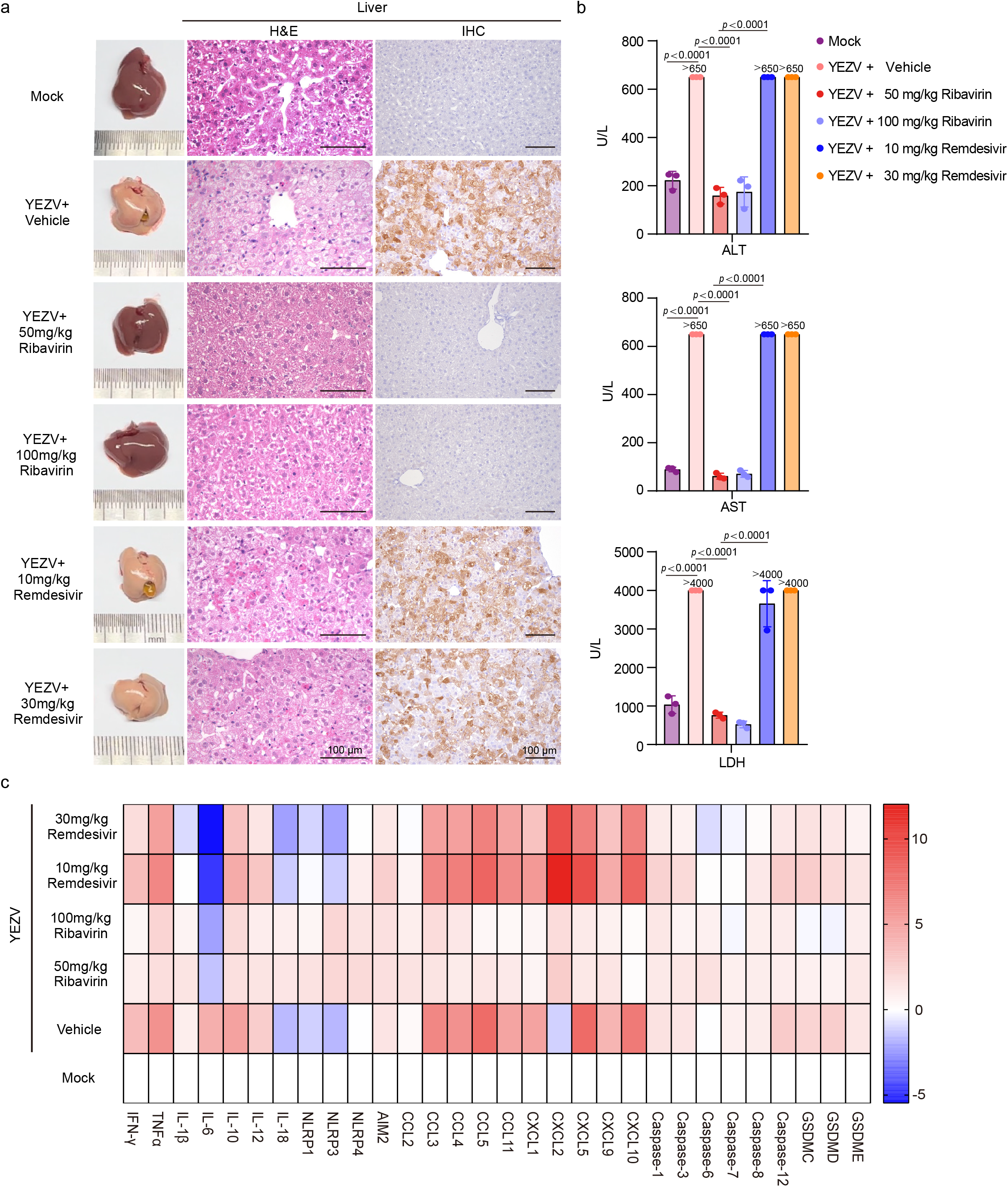
Ribavirin treatment mitigates MA-YEZV-induced hepatic pathology and inflammation. Three-week-old male C57BL/6J mice were intraperitoneally inoculated with 1 TCID_50_ of MA-YEZV. Ten days post-infection, liver and blood samples were collected for pathological, biochemical, and cytokine analyses. (a) Liver histopathology assessment, including gross morphology, hematoxylin and eosin (H&E) staining, and immunohistochemistry (IHC) staining using an antibody against the YEZV nucleoprotein (NP), comparing mock-infected mice to MA-YEZV-infected mice treated with ribavirin or remdesivir. Scale bars, 100 μm (H&E), 50 μm (IHC). (b) Serum biochemical markers are indicative of liver injury. Data are presented as mean ± standard deviation (n = 3 biological replicates). ALT, alanine aminotransferase; AST, aspartate aminotransferase; LDH, lactate dehydrogenase. Statistical significance was determined by one-way ANOVA (two-sided) followed by Tukey’s multiple-comparison test to adjust for multiple testing. Exact *p* values are provided in the figure legends. (c) Hepatic cytokine mRNA expression levels measured by RT-qPCR in infected mice treated with ribavirin and remdesivir (n = 3 biological replicates, relative expression). Source data are provided as a Source Data file.

Parallel assessment of the spleen showed extensive white pulp necrosis, enhanced macrophage phagocytosis, and marked histiocytic proliferation in both red and white pulp of vehicle controls. These pathological features were significantly mitigated by ribavirin treatment, with only mild histiocytic proliferation restricted to the red pulp (Supplementary Fig. 11). Testes from vehicle controls displayed structural damage in seminiferous tubules, including apoptotic bodies, reduced germ cells, and a residual single layer of spermatogonia with sparse Sertoli cells. Treatment with 50 mg/kg ribavirin markedly alleviated this damage, causing only mild interstitial fibrosis. Notably, 10 mg/kg remdesivir, despite showing no clinical benefit, completely reversed MA-YEZV-induced pathological lesions. Conversely, high-dose ribavirin (100 mg/kg) and remdesivir (30 mg/kg) reduced germ cell counts in specific seminiferous tubules, suggesting potential reproductive toxicity (Supplementary Fig. 11). Remdesivir-treated mice showed severe pathology in the spleen similar to vehicle controls, with no significant improvement (Supplementary Fig. 11).

Consistent with the histological findings, ribavirin treatment normalized infection-induced hematological and biochemical abnormalities. Levels of ALT, AST, LDH, TBA, ALP, and total bilirubin were restored to baseline, confirming protection against hepatic injury (Fig. 6b, Supplementary Fig. 12, Supplementary Table 9). Additionally, serum ALB, GLB, and TP levels were replenished, indicating recovery of hepatic synthetic function. Hematological parameters showed complete reversal of infection-driven reductions in white blood cells, lymphocytes, and platelets. In contrast, remdesivir failed to significantly improve these parameters (Fig. 6b, Supplementary Fig. 12, Supplementary Table 10).

Ribavirin treatment also reversed the elevated levels of proinflammatory mediators (IL-6, IL-10, IFN-γ, MCP-1, TNF-α) in serum of MA-YEZV-infected mice, restoring them to near-baseline levels (Supplementary Fig. 13). Although remdesivir was clinically and virologically ineffective, it paradoxically reduced IL-6, IL-10, IFN-γ, and TNF-α levels. At the transcriptional level, ribavirin broadly suppressed the infection-induced upregulation of hepatic cytokines, with the most pronounced effect on *IL-6*. While remdesivir enhanced *IL-6* suppression, it significantly upregulated *CXCL2*, a chemokine that recruits neutrophils to the inflamed liver via CXCR2 binding (Fig. 6c).^24^ Collectively, these results demonstrate that ribavirin treatment effectively mitigates pathological damage and ameliorates inflammatory responses.

## Discussion

The development of animal models that faithfully recapitulate human disease is critical for understanding the pathogenesis of emerging viruses and evaluating potential therapeutics. In this study, we successfully generated a mouse-adapted strain of Yezo virus (MA-YEZV) capable of causing lethal infection in immunocompetent mice, addressing a significant gap in the study of orthonairoviruses. This model not only mirrors key clinical features of human YEZV infection, but also provides a robust platform for investigating viral pathogenesis and antiviral interventions.

Our serial passaging of YEZV in C57BL/6J mice yielded MA-YEZV, a strain exhibiting enhanced replication kinetics *in vitro* and lethal pathogenicity *in vivo*. The adapted strain induced hallmark symptoms of human YEZV infection, including thrombocytopenia, leukopenia, and elevated liver transaminases, with hepatic injury as the predominant pathology^11,15,16^. These findings align with clinical reports of YEZV-infected patients, underscoring the translational relevance of this model.

Previous research established that that the human-derived YEZV strain HH003-2019 causes only a limited, non-lethal infection in immunocompetent mice, a pathogenic profile similar to that of the parental strain T-HLJ01 used here^18^. Consequently, studies have relied on immunodeficient AG129 mice to model severe YEZV induced acute hepatitis, which recapitulates key biochemical markers of liver injury, including ALT, AST, ALB, GLB and TP^18^. In contrast, the mouse-adapted MA-YEZV strain developed in this study induces lethal hepatitis in immunocompetent C57BL/6J mice while preserving the intrinsic hepatotropism of YEZV. Moreover, MA-YEZV infection elicits marked leukopenia and thrombocytopenia, the hallmark clinical features of human YEZV infection that are not observed in the AG129 mouse models^18^. These findings demonstrate that the MA-YEZV immunocompetent mouse model not only captures the hepatic tropism and severe pathology of YEZV but also more faithfully reproduces the immune-mediated hematological abnormalities seen in patients. This provided a more physiologically relevant platform for investigating virus–host interactions and evaluating potential antiviral interventions.

The liver tropism observed in MA-YEZV-infected mice is consistent with the hepatic involvement seen in other orthonairovirus infections, such as Crimean-Congo hemorrhagic fever virus (CCHFV)^25^. Notably, MA-YEZV targeted multiple hepatic cell types, including hepatocytes, hepatic stellate cells, and liver sinusoidal endothelial cells, suggesting a broad cellular tropism that may contribute to its pathogenicity. The robust inflammatory response, characterized by elevated proinflammatory cytokines (e.g., IFN-γ, TNF-α, IL-6) and inflammasome activation, likely exacerbates tissue damage and disease severity. This hyperinflammatory state mirrors the cytokine storms reported in severe human cases of orthonairovirus infections, highlighting the utility of MA-YEZV for studying immune-mediated pathology^21^.

While there is no current evidence of human-to-human YEZV transmission or detection of infectious virus in clinical specimens, our MA-YEZV infection model demonstrates that the virus can replicate to high titers in multiple tissues and be shed into body fluids. Despite this, neither horizontal nor vertical transmission occurred among co-housed or breeding mice, suggesting that efficient spread may be limited by specific host or environmental factors. Notably, intranasal inoculation in mice established a systemic infection, confirming that the respiratory tract can serve as a viable portal of entry. These observations indicate that YEZV possesses the fundamental capacity for replication and shedding in a mammalian host, yet its natural transmission likely depends on conditions not fully met in our experimental setting. These findings underscore the need for ongoing surveillance to better understand YEZV transmission dynamics and its potential for adaptation.

Genomic analysis revealed 31 non-synonymous mutations in MA-YEZV, distributed across viral proteins critical for replication (RdRp), immune evasion (OTU protease), and entry (Gc glycoprotein). Of these, 10 mutations were enriched in human-derived YEZV strains, suggesting they may represent adaptive changes favoring mammalian host infection. The mutations in the OTU domain, known to interfere with host antiviral responses, and those in the Gc glycoprotein, which may alter receptor binding, are particularly compelling candidates for future mechanistic studies^26,27^. The synergistic effects of these mutations likely underpin the enhanced virulence of MA-YEZV, as observed in other adapted viruses.

Ribavirin demonstrated remarkable efficacy against MA-YEZV, reducing viral loads, mitigating pathology, and improving survival. Its success contrasts with the inefficacy of remdesivir, suggesting that YEZV’s RdRp may preferentially incorporate ribavirin’s guanosine analog over remdesivir’s adenosine analog. This finding has immediate implications for clinical management, as ribavirin has been shown to be effective against other tick-borne viruses^28^. However, the potential reproductive toxicity observed at high doses warrants caution and further investigation.

While MA-YEZV recapitulates many features of human YEZV infection, the model has limitations. The absence of neurological pathology, despite clinical reports of neurological symptoms in patients, suggests that alternative models (e.g., non-human primates) may be needed to fully capture the disease spectrum^16^. Additionally, the lack of horizontal or vertical transmission in mice contrasts with the tick-borne spread of YEZV in nature, highlighting the need for vector-mediated transmission studies. Future work should focus on: i) Mechanistic studies to link specific mutations in MA-YEZV to its enhanced virulence and tropism; ii) Therapeutic development, including combination therapies to mitigate ribavirin’s side effects and novel antivirals targeting YEZV’s unique RdRp features; iii) Transmission models incorporating tick vectors to better mimic natural infection dynamics.

The MA-YEZV model represents a significant advance in the study of emerging orthonairoviruses, providing a tractable system to investigate pathogenesis, host responses, and antiviral strategies. Its alignment with human disease features underscores its potential to accelerate the development of countermeasures against YEZV and related viruses. This work also highlights the importance of adaptive evolution in viral emergence and the need for vigilant surveillance of tick-borne pathogens with zoonotic potential.

## Methods

### Animal ethics and biosafety

All animal procedures were reviewed and approved by the Animal Experiment Committee of Laboratory Animal Center in Jilin University (approval number: 11968). Studies were carried out in strict accordance with the recommendations in the Guide for the Care and Use of Laboratory Animals.

The generation and handling of the mouse-adapted virulent strain (MA-YEZV) were conducted under explicit approval and oversight from the Institutional Biosafety Committee (IBC) of Jilin University. The IBC reviewed and approved the protocol for creating an enhanced pathogen, confirming its compliance with national biosafety regulations and institutional policies for the oversight of dual-use life sciences research. All work with infectious YEZV, including the parental T-HLJ01 and mouse-adapted MA-YEZV strains, was performed under Biosafety Level 3 (BSL-3) containment.

### Cells

Vero (SCSP-520), HEK-293T (SCSP-502), A549 (SCSP-503), and Hep G2 (SCSP-510) cells were obtained from the National Collection of Authenticated Cell Cultures (Shanghai, China). Vero, HEK 293T, A549 cells were cultured in Dulbecco’s Modified Eagle Medium (DMEM, Gibco, USA) supplemented with 10% (v/v) fetal bovine serum (FBS; Gibco) and 1% (v/v) penicillin-streptomycin (Gibco). Hep G2 cells were cultured in Minimum Essential Medium (MEM, Gibco, USA) supplemented with 10% (v/v) fetal bovine serum (FBS; Gibco), 1% non-essential amino acids (Gibco), and 1% (v/v) penicillin-streptomycin (Gibco). Cells were maintained at 37°C in a humidified 5% CO_2_ atmosphere and confirmed to be mycoplasma-free prior to use.

### Antibodies

An antibody against the Yezo virus nucleoprotein (NP) was generated by immunizing rabbits with recombinant NP antigen (Detai Bioengineering, Nanjing, China). The recombinant protein was produced by cloning a codon-optimized YEZV-NP gene into pET-30 expression vector. Following expression in *Escherichia coli*, the protein was purified using affinity chromatography. The purified NP protein was then used to immunize New Zealand White rabbits three times. Antisera were collected and subsequently affinity-purified to obtain the final anti-NP polyclonal antibody.

The commercial primary antibodies used in this study were as follows: anti-albumin (AF3329), anti-CD31 (AF3628), and anti-desmin (AF3844) (R&D Systems, USA); anti-F4/80 (ab320060) (Abcam, USA); anti-NLRP3 (15101S), anti-beta-Actin (4970S), and anti-GAPDH (2118S) (Cell Signaling Technology, USA).

### Mice

Specific pathogen-free (SPF) three-week-old BALB/c, C57BL/6J, and Kunming mice were obtained from Changsheng Biotechnology (Benxi, China). Mice were housed at 22–25°C with 40–70% relative humidity under a 12-hour light/dark cycle (lights on 07:00 –19:00) in individually ventilated cages, with unrestricted access to food and water.

### Virus isolation

The YEZV strain T-HLJ01 (GenBank accession numbers ON563268–ON563270) was isolated from tick-positive samples identified in our previous study^15^. Briefly, tick homogenates were centrifuged, filtered, diluted to 1:20, and inoculated onto Vero cells. After 2 hours incubation at 37°C, cells were washed and fresh medium added. Following three blind passages, supernatants were collected and tested using RT-qPCR.

The MA-YEZV strain was isolated from mouse liver tissue at passage 40 (P40). Liver tissues were homogenized in 500 μL DMEM per 50 mg of tissue, centrifuged, and the supernatant diluted 1:50 to inoculate Vero cells.

### Tissue passaging

Three-week-old female C57BL/6J mice were intraperitoneally inoculated with 10^4^ TCID_50_ of YEZV strain T-HLJ01 (passage 0, P0). At 3 days post-infection (dpi), mice were euthanized, and liver tissues were collected. For every 50 mg liver tissue, 500 μL DMEM was added, and tissues homogenized at 70 Hz for 90 seconds. Homogenates underwent two freeze-thaw cycles to release viral particles, followed by centrifugation at 12,900 g for 30 minutes at 4°C. The supernatant was injected intraperitoneally into naïve C57BL/6J mice (passage 1, P1). Serial passages were continued similarly. Mice were monitored daily for clinical signs, with pathological changes recorded at necropsy. Due to mortality observed at P30, the passage interval was shortened to 2.5 days. Passaging continued until P40, at which point YEZV was isolated from P40 mouse liver tissues using Vero cells. To ensure enhanced safety, all murine infection experiments involving YEZV were performed under Biosafety Level 3 (BSL3) containment conditions.

### Viral genome amplification

Viral RNA was extracted from 140 μL of culture supernatant, mouse sera, tissue homogenates, or swabs using the TIANamp Virus RNA kit (TIANGEN, China)^29^. cDNA synthesis was performed with the PrimerScript™ RT reagent Kit with gDNA Eraser (TaKaRa, China).^30^ Semi-nested PCR reactions were performed in a total volume of 25 μL, containing 12.5 μL Premix Taq (TaKaRa, China), 1 μL of cDNA, and 10 pmol of primers (Supplementary Table 11).^31^ Cycling conditions were as follows: initial denaturation at 94°C for 5 minutes; 35 cycles of 94°C for 30 seconds, 50°C for 30 seconds, and 72°C for 30–60 seconds (depending on the amplicon length); followed by a final extension at 72°C for 5 minutes. The second-round PCR used the first-round products as templates, with the same reaction composition and cycling parameters as the first round. PCR products were sequenced bidirectionally using the Sanger method.

### RT-qPCR

Viral RNA extraction and reverse transcription were performed as previously described. For virus quantification, YEZV-specific primers and probes are listed in Supplementary Table 11. Quantitative PCR was conducted using the Premix Ex Taq™ kit (Takara, Japan) on the StepOne Plus™ Real-Time System (Thermo Fisher Scientific, USA)^32^. A cycle threshold (CT) value less than 40 was considered positive. Standard curves (y = -3.356x + 41.53) converted CT values to absolute viral copy numbers.

### Virus titration assay

Supernatants from infected cell cultures or tissue homogenates were serially diluted 10-fold from 10^−1^ to 10^−8^. For each dilution, 100 μL viral suspension was added to eight replicate wells of a 96-well plate seeded with 1 × 10^4^ vero cells per well. Cytopathic effects (CPE) were monitored daily for 5 days. Infection was assessed by indirect immunofluorescence staining using anti-YEZV NP. The 50% tissue culture infectious dose (TCID_50_) was calculated using the Reed–Muench method^33,34^.

### Mouse infection experiments

To evaluate MA-YEZV pathogenicity, three-week-old C57BL/6J mice were intraperitoneally inoculated with 10^4^, 10^2^, or 1 TCID_50_ of MA-YEZV. For route comparison, age-matched male C57BL/6J mice were inoculated with 1 TCID_50_ via intranasal (i.n.) or subcutaneous (s.c.) routes. Strain susceptibility was evaluated in BALB/c and Kunming mice following i.p. injection of 1 TCID_50_ MA-YEZV. Clinical signs and mortality were recorded daily. Tissue, serum, and swab samples were collected for viral load quantification by RT-qPCR. For horizontal transmission studies, three-week-old male C57BL/6J mice (index group; n = 3) were intraperitoneally inoculated with 1 TCID_50_ of MA-YEZV. After 3 days, individual index mice were co-housed with three naïve male contacts. Clinical signs and mortality were monitored for 14 days. Tissue and serum samples from both groups were collected at 3, 7, 10, and 14 dpi for viral load quantification via RT-qPCR.

For vertical transmission studies, 10-week-old pregnant C57BL/6J female mice (n = 3) at embryonic day 12.5 were intraperitoneally inoculated with 1 TCID_50_ of MA-YEZV. After delivery, viral loads in tissues and serum samples of 1-day-old newborn mice were assessed by RT-qPCR.

### Antiviral efficacy against MA-YEZV infection

Ribavirin (HY-B0434), remdesivir (GS-5734) and its active metaboliteGS-441524 were obtained from MedChemExpress (USA) and dissolved in vehicle (5% DMSO, 40% PEG300, 5% Tween-80, and 50% saline)^35,36^. Three-week-old male C57BL/6J mice were intraperitoneally infected with 1 TCID_50_ of MA-YEZV. At 2 hours post-infection (hpi), mice received daily intraperitoneal injections of ribavirin (50 or 100 mg/kg), remdesivir (GS-441524, 25 or 50 mg/kg) and its active metabolite GS-5734 (10 or 30 mg/kg) for 8 days. Infection controls and mock groups received equivalent volumes of vehicle. Clinical manifestations and survival rates were recorded for 14 dpi. Tissue and serum samples were collected at 3, 7, and 10 dpi for viral load quantification, histopathology, and hematological analysis.

For the *in vitro* antiviral assays, Vero cells were infected with MA-YEZV at an MOI of 0.01. Following a 2-hour adsorption period, the inoculum was replaced with maintenance medium containing serial dilutions of ribavirin (HY-B0434; 0.01–200 μM), remdesivir (GS-441524; 0.01– 200 μM), or its active metabolite GS-5734 (0.01–200 μM). Supernatants were harvested at 5 dpi for viral titration, and the half-maximal inhibitory concentration (IC_50_) for each compound was calculated by nonlinear regression using GraphPad Prism 9. The potential cytotoxicity of each drug concentrations was assessed in parallel using a CCK-8 assay^37^.

### Hematoxylin and eosin staining and immunohistochemistry

Hematoxylin and eosin (H&E) staining was performed as previously described^38^. Tissue sections were stained with hematoxylin for 5 minutes, then rinsed under running water for 5 minutes to develop a blue nuclear background. Sections were rapidly differentiated in 1% acid alcohol, with differentiation halted by rinsing under running water (guided by microscopic examination). After dehydration and clearing, sections were examined under a light microscope (Zeiss, Germany) to assess tissue damage.

Tissue sections were subjected to antigen retrieval by heating in citrate buffer (10 mM, pH 6.0) at 95°C for 20 minutes. Endogenous peroxidase activity was blocked with 3% hydrogen peroxide in methanol at room temperature for 10 minutes. After washing with PBS, sections were blocked with 5% goat serum for 30 minutes to reduce nonspecific binding. Then, sections were incubated overnight at 4°C with the primary antibody anti-YEZV NP (diluted 1:200). Following washing, sections were incubated with HRP-conjugated secondary antibody at room temperature for 1 hour. Visualization was performed using 3,3’-diaminobenzidine (DAB), followed by counterstaining with hematoxylin. After dehydration, clearing, and mounting, the sections were examined under a light microscope (Zeiss, Germany)^39^.

### Immunofluorescence staining

Immunofluorescence (IF) staining was performed on YEZV-infected Vero cells and paraffin-embedded tissue sections using a standardized protocol^40,41^. Cells were fixed with 4% paraformaldehyde for 30 minutes and permeabilized with 1% Triton X-100 for 10 minutes. For tissue sections, endogenous peroxidase activity was quenched with 3% hydrogen peroxide for 10 minutes at room temperature. All specimens were blocked with 5% bovine serum albumin (BSA) for 1 hour at room temperature, then incubated overnight at 4°C with anti-YEZV nucleoprotein (NP) antibody (1:200 in 1% BSA-PBS). After washing, samples were incubated in the dark at 37°C for 1 hour with FITC-conjugated goat anti-rabbit IgG (H+L) secondary antibody (1:500). Nuclei were counterstained with DAPI, and fluorescence images acquired using an inverted fluorescence microscope (Zeiss, Germany).

For multiplex immunofluorescence staining of mouse liver tissues, co-staining was performed for YEZV and specific markers: albmin (hepatocytes), CD31 (liver sinusoidal endothelial cells), F4/80 (Kupffer cells), and desmin (hepatic stellate cells). Nuclei were stained with DAPI, and images captured using the Zeiss inverted fluorescence microscope.

### Cytokine analysis

Cytokine levels in mouse serum were quantified using the BD™ Cytometric Bead Array (CBA) Mouse Inflammation Kit (BD Biosciences, USA)^42^. For each assay tube, 50 μL diluted standard or mouse serum sample was combined with 50 μL mixed capture beads specific to each cytokine. Then 50 μL PE-conjugated Detection Reagent was added, and samples were incubated at room temperature in the dark for 2 hours. After adding 1 mL wash buffer, samples were centrifuged at 200×g for 5 minutes, supernatant discarded, and beads were resuspended in 300 μL wash buffer. Samples were acquired by flow cytometry, and fluorescence intensities for each cytokine were recorded. Data analysis was performed using FCAP Array™ software (BD Biosciences) by gating bead populations, generating standard curves, and determining mean fluorescence intensity of each cytokine. Cytokine concentrations were calculated from the corresponding standard curves.

To analyze hepatic transcription, total RNA was extracted using RNA-easy Isolation Reagent (Vazyme, China)^43^. Reverse transcription was performed using the PrimeScript^™^ RT reagent Kit with gDNA Eraser (TaKaRa, Japan). Quantitative PCR was carried out using TB Green^®^ Premix Ex Taq™ II (TaKaRa, Japan) in a 20 μL reaction containing 10 μL 2× TB Green Premix, primer at 0.4 μM (Supplementary Table 11), 10–100 ng cDNA, 0.4 μL ROX Reference Dye, and nuclease-free water. Reactions were run in technical triplicate, with no-template controls to detect contamination. ACTB was used as the internal reference gene, and relative gene expression was calculated using the 2 ^−ΔΔCt^ method^44^.

For cytokine protein quantification, total protein was extracted from mouse liver tissues using pre-chilled strong RIPA lysis buffer (Beyotime, China) supplemented with 1% PMSF for cytokine protein quantification^45^. Approximately 20 mg tissue per sample was homogenized in 200 μL lysis buffer at 70 HZ for 90 seconds, then centrifuged at 12,900 g for 15 minutes at 4°C. Supernatant was mixed with loading buffer and boiled for 10 minutes for denaturation. Proteins were separated by SDS–PAGE and transferred onto PVDF membranes, which were blocked with 5% non-fat milk in TBST for 1 hour at room temperature. Membranes were incubated overnight at 4°C with primary antibodies (YEZV-NP, 1:1000; NLRP3, 1:1000; beta-Actin, 1:5000; GAPDH, 1:5000) in blocking buffer. After washing, membranes were incubated with HRP-conjugated secondary antibodies for 1 hour at room temperature. Protein bands were visualized using enhanced chemiluminescence and captured with an imaging system. Band intensities were quantified using ImageJ software, with relative expression levels calculated accordingly.

### Statistics and Reproducibility

Statistical analyses were conducted using GraphPad Prism 9. The Shapiro–Wilk test was used to assess data normality. Normally distributed data are presented as the mean ± standard deviation (SD). For multiple group comparisons, one-way analysis of variance (ANOVA) was performed, followed by Tukey’s post hoc test as appropriate. For two-group comparisons, an unpaired two-tailed Student’s *t*-test was used; the Mann–Whitney U test was used for non-normally distributed data. All statistical tests were two-tailed. A *p*-value < 0.05 was considered statistically significant, with significance annotated in figures.

## Data Availability

The YEZV strain T-HLJ01 (GenBank accession numbers ON563268–ON563270) used in this study was obtained from previously reported isolates (https://www.ncbi.nlm.nih.gov/nuccore/ON563268.1; https://www.ncbi.nlm.nih.gov/nuccore/ON563269.1; https://www.ncbi.nlm.nih.gov/nuccore/ON563270.1). The mouse-adapted Yezo virus strain generated in this study (Yezo virus strain MA-HLJ01-p40) has been deposited in the GenBank database under accession numbers PV976754-PV976756 (https://www.ncbi.nlm.nih.gov/nuccore/PV976754.2; https://www.ncbi.nlm.nih.gov/nuccore/PV976755.2; https://www.ncbi.nlm.nih.gov/nuccore/PV976756.2).

## Acknowledgements

The study was funded by the National Key Research and Development Program of China (2024YFD1800103, Q.L.) and the National Natural Science Foundation of China (82341105 and 82372250, Q.L.).

## Author Contributions Statement

W.X., M.P., Q.T., F.J., L.S., and Y.Z. performed experiments and analyzed data. N.L., Y.W., and Q.L. designed and supervised the project. W.X., N.L., and Q.L. interpreted the results. W.X. wrote the original manuscript. N.L. and Q.L. revised the manuscript. All authors reviewed and proofread the manuscript.

