## Supplementary Figures 1 to 13 and Supplementary Tables 1 to 11 for "A mouse-adapted Yezo virus model for antiviral testing in immunocompetent mice"

**This PDF file includes:**

Supplementary Figures 1 to 13

Supplementary Tables 1 to 11

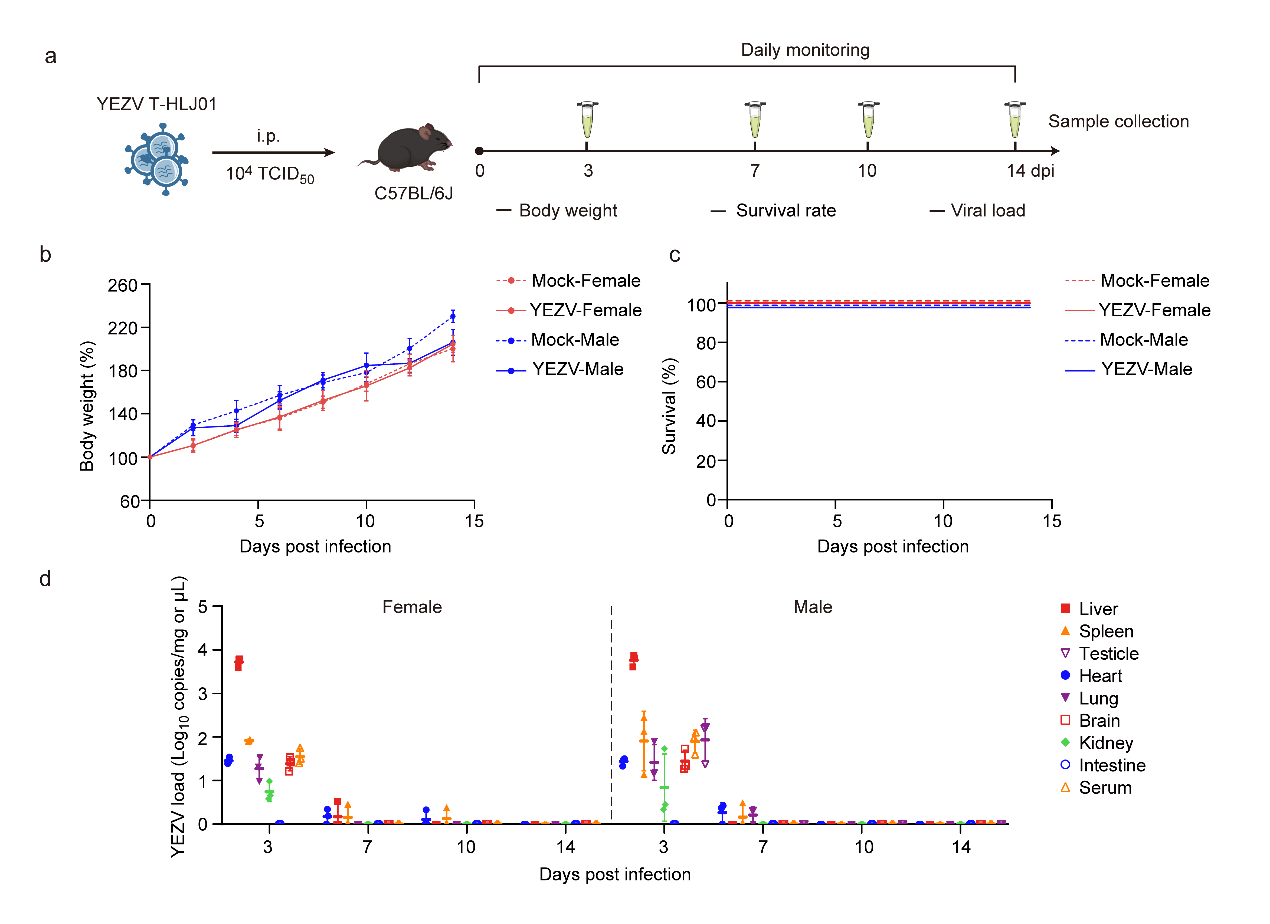
**Supplementary Fig. 1 The YEZV strain T-HLJ01 establishes a transient subclinical infection in C57BL/6J mice**

(a) Schematic of the experimental timeline. Three-week-old female and male C57BL/6J mice were intraperitoneally (i.p.) inoculated with 10^4^ TCID₅₀ of YEZV strain T-HLJ01. Body weight and survival were monitored daily. Tissues and serum were collected at indicated time points for viral load analysis.

(b, c) Relative body weight change (b) and survival (c) of T-HLJ01-infected C57BL/6J mice (n=6 independent mice per sex) compared to mock-infected controls (n=6 independent mice per sex).

(d) Viral loads in tissues and serum of T-HLJ01-infected mice at 3, 7, 10 and 14 days post-infection, quantified by RT-qPCR (n=3 biological replicates).

Data in (b) and (d) are presented as mean ± standard deviation (SD). Source data are provided as a Source Data file.

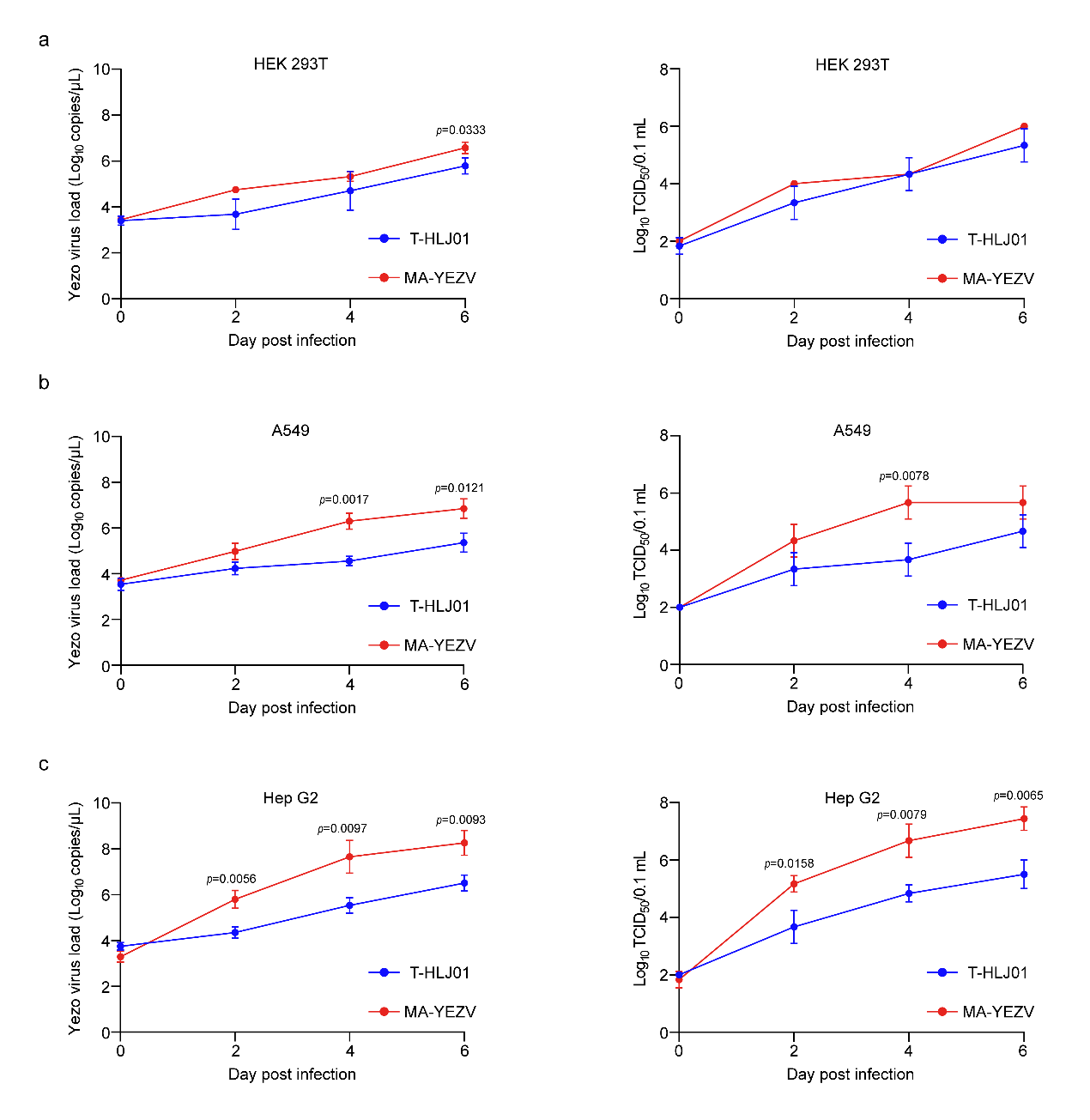
**Supplementary Fig. 2 MA-YEZV exhibits enhanced replication in human cell lines compared to the parental T-HLJ01 strain**

(a–c) Viral replication kinetics of T-HLJ01 and MA-YEZV in human embryonic kidney (HEK 293T; a), lung carcinoma (A549; b), and hepatocellular carcinoma (Hep G2; c) cells. Viral RNA copy numbers (left) and infectious titers (right) in cell culture supernatants were measured over 7 days. Data are presented as mean ± standard deviation (n=3 biological replicates). Statistical significance between two groups at each time point was determined using a two-sided unpaired Student’s t-test. Exact *p* values are indicated in the figure. Source data are provided as a Source Data file.

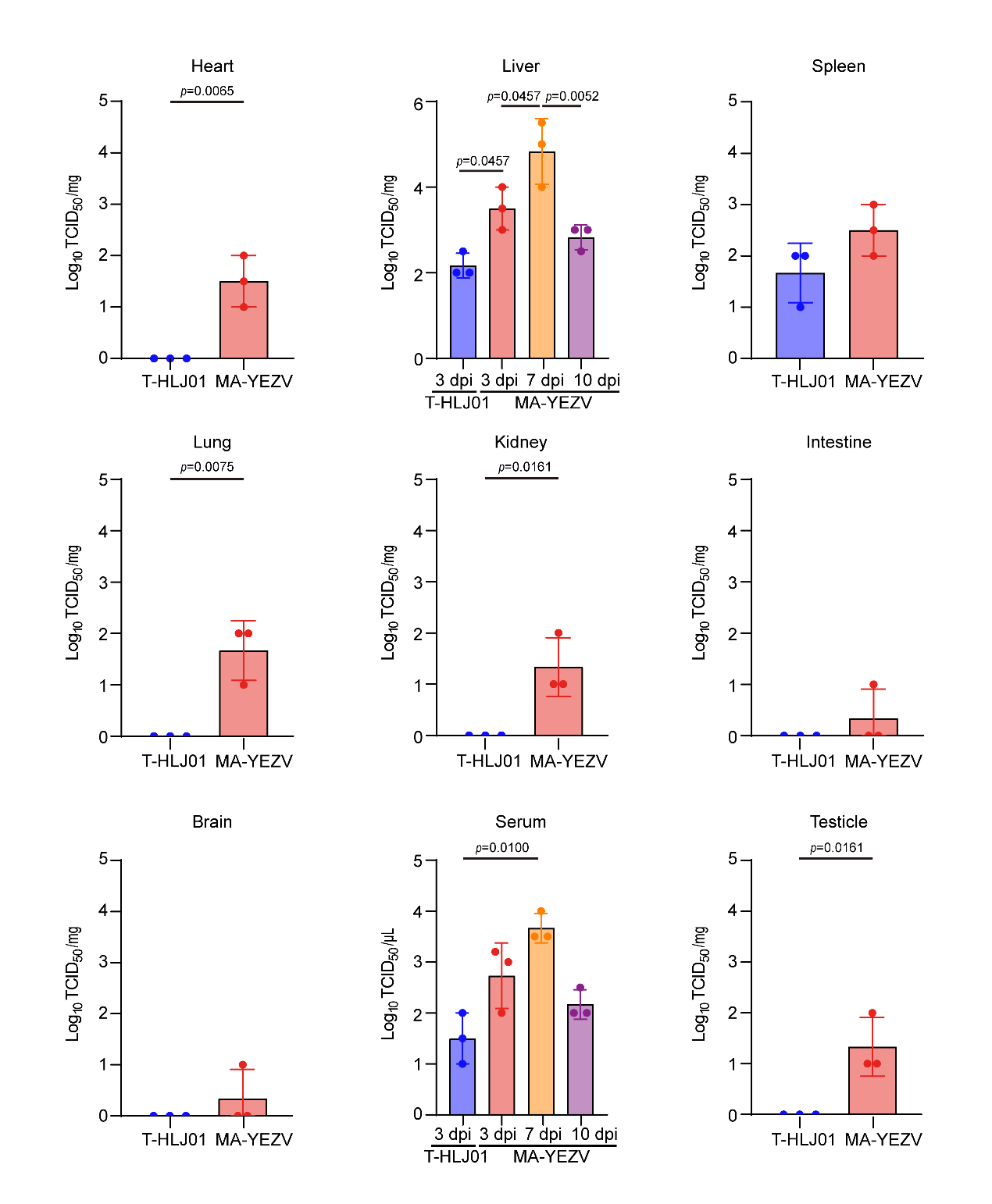
**Supplementary Fig. 3 Comparison of tissue viral titers between mice infected with T-HLJ01 and MA-YEZV**

Viral titers (TCID_50_/mg tissue or µL serum) were measured at 3 dpi in mice intraperitoneally inoculated with 10^4^ TCID_50_ of T-HLJ01 or 1 TCID_50_ of MA-YEZV. The liver was the primary site of replication for both strains. MA-YEZV achieved significantly higher titers in most tissues and serum compared to T-HLJ01, demonstrating enhanced systemic dissemination. Data are presented as mean ± SD (n=3 biological replicates). Statistical significance between two groups at each time point was determined using a two-sided unpaired Student’s *t*-test. For comparisons among multiple groups, one-way ANOVA (two-sided) was performed, followed by Tukey’s multiple-comparison test to adjust for multiple testing. Exact *p* values are indicated in the figures. Source data are provided as a Source Data file.

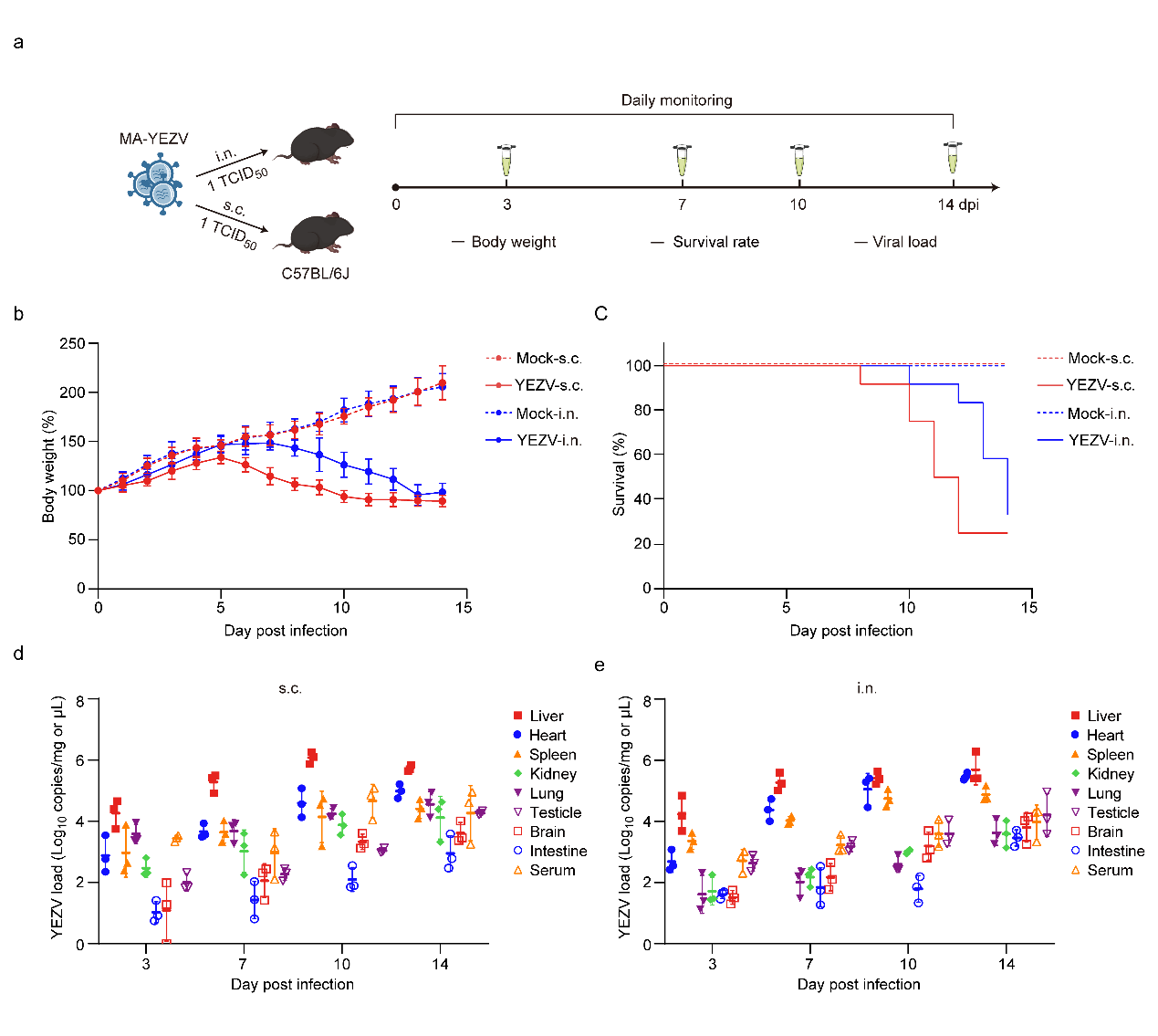

**Supplementary Fig. 4 MA-YEZV is lethal in C57BL/6J mice via subcutaneous and intranasal inoculation routes.**

(a) Experimental timeline. Three-week-old male C57BL/6J mice were inoculated with 1 TCID_50_ of MA-YEZV via subcutaneous (s.c.) or intranasal (i.n.) routes. Body weight and survival were monitored daily; tissues and serum were collected at indicated time points for analysis.

(b, c) Relative body weight changes (b) and survival (c) of mice infected via s.c. or i.n. routes (n=12 independent mice per group) compared to mock-infected controls (n=6 independent mice).

(d, e) Tissue and serum viral loads, measured by RT-qPCR at 3, 7, 10, and 14 days post-infection following s.c. (d) or i.n. (e) MA-YEZV inoculation (n=3 biological replicates).

Data in (b, d, and e) are presented as mean ± standard deviation. Source data are provided as a Source Data file.

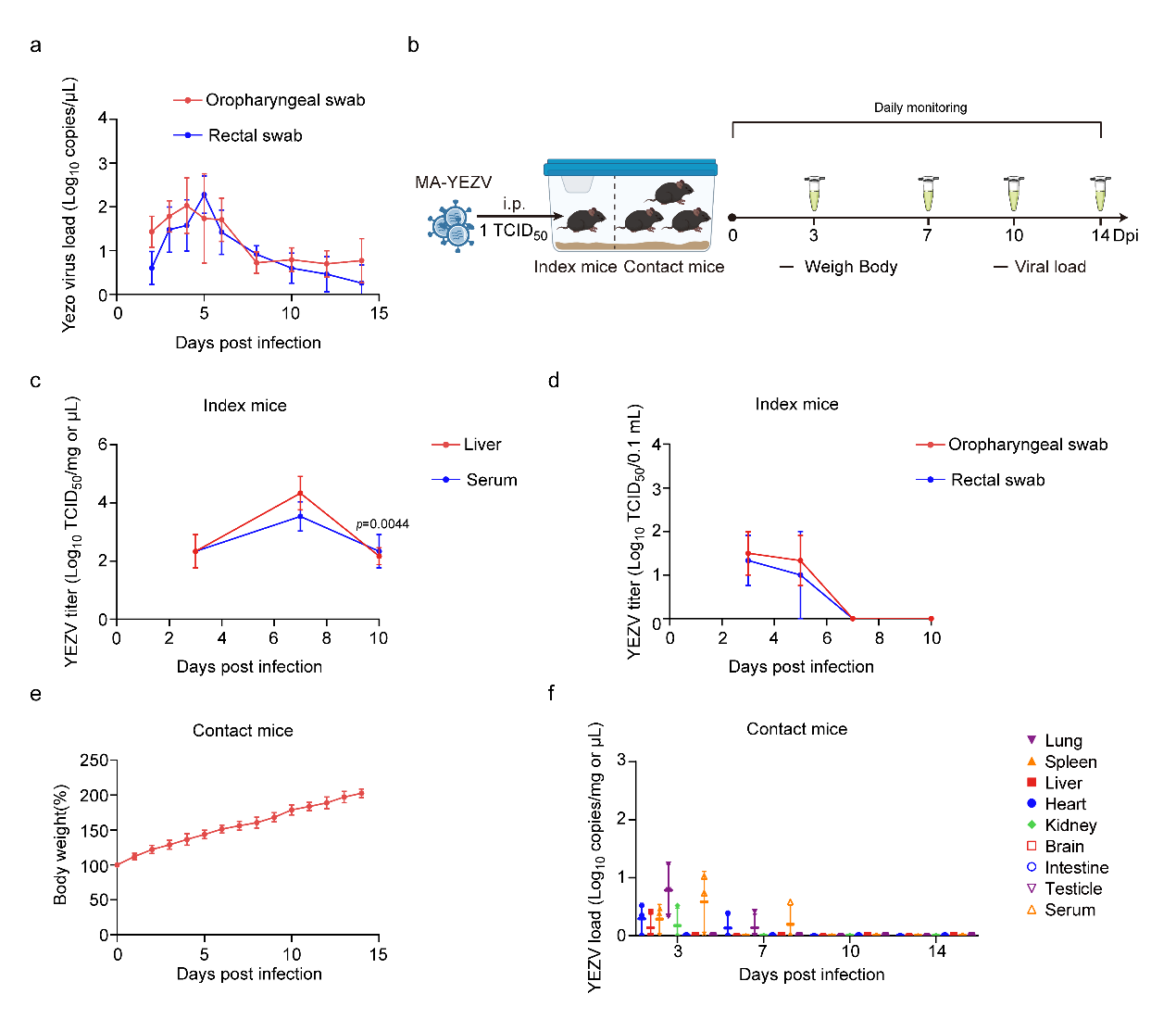
**Supplementary Fig. 5 Assessment of MA-YEZV shedding and horizontal transmission in mice**

(a) Viral RNA kinetics in oropharyngeal and rectal swabs from intraperitoneally (i.p.) infected mice (n=3 biological replicates), measured by RT-qPCR over 14 days.

(b) Experimental design for contact transmission. Mice were infected (1 TCID₅₀, i.p.) and housed individually for 3 days before co-housing with naïve mice (3:1 ratio).

(c, d) Infectious viral titers in the liver and serum (c), and in rectal and oropharyngeal swabs (d) of index mice at 3, 7, and 10 days post-infection (n=3 biological replicates). Statistical significance between two groups at each time point was determined using a two-sided unpaired Student’s t-test. Exact *p* values are indicated in the figures.

(e) Body weight change of contact mice (n=9 independent mice) monitored over 15 days of co-housing.

(f) Viral loads in tissues and serum of contact mice at 3, 7, 10, and 14 days post co-housing, measured by RT-qPCR (n=3 biological replicates).

Data are presented as mean ± standard deviation (a, c, d, e and f). Source data are provided as a Source Data file.

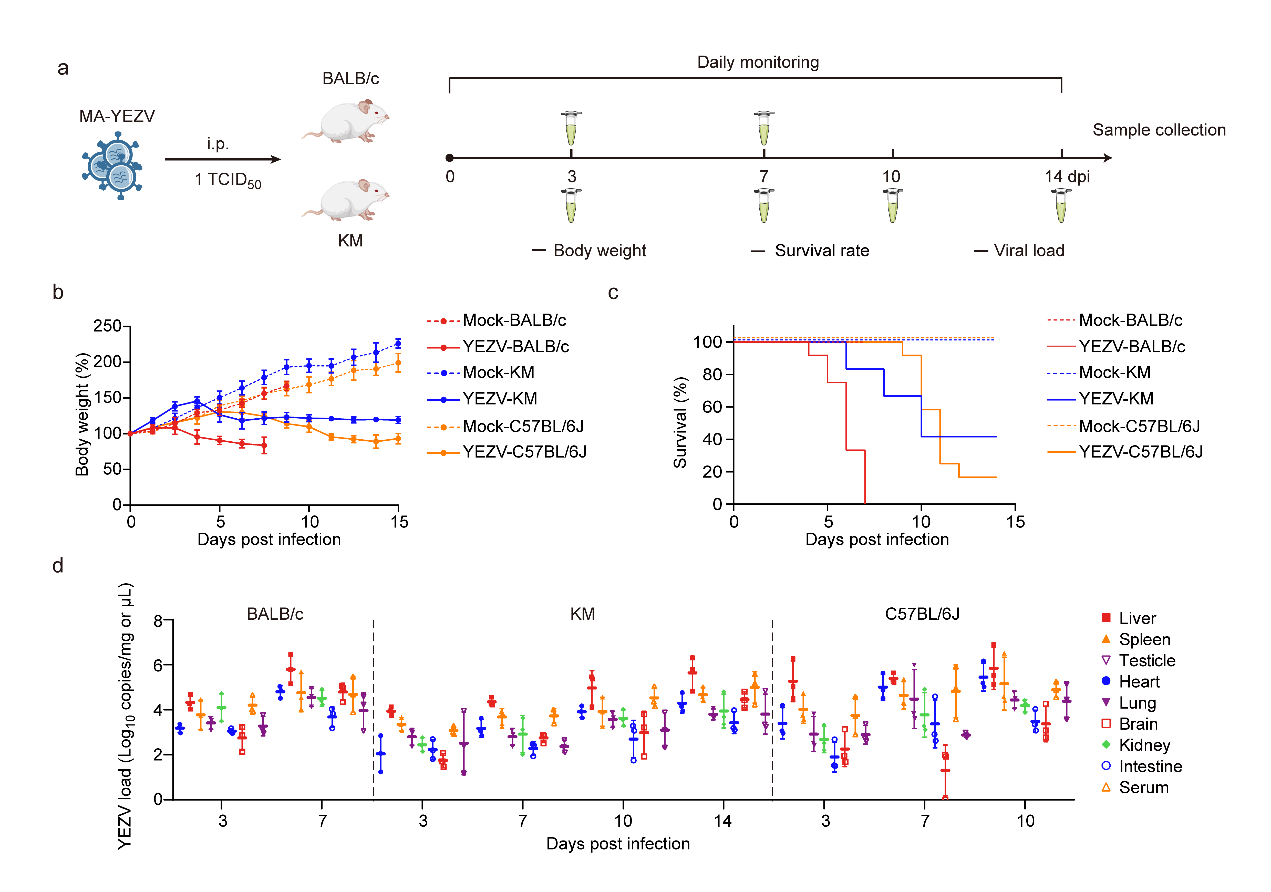
**Supplementary Fig. 6 Lethal infection by MA-YEZV in multiple immunocompetent mouse strains.**

(a) Experimental timeline. Three-week-old male BALB/c and Kunming (KM) mice were intraperitoneally (i.p.) inoculated with 1 TCID₅₀ of MA-YEZV. For comparison, data from C57BL/6J mice (from Fig. 2) are included.

(b, c) Clinical course showing relative body weight change (b) and survival (c) in infected mice (n=12 independent mice per strain) versus mock-infected controls (n=6 independent mice).

(d) Viral loads in tissues (heart, liver, spleen, lung, kidney, intestine, brain) and serum of infected BALB/c and KM mice at 3, 7, 10, and 14 dpi, measured by RT-qPCR (n=3 biological replicates).

Data are presented as mean ± standard deviation (b and d). Source data are provided as a Source Data file.

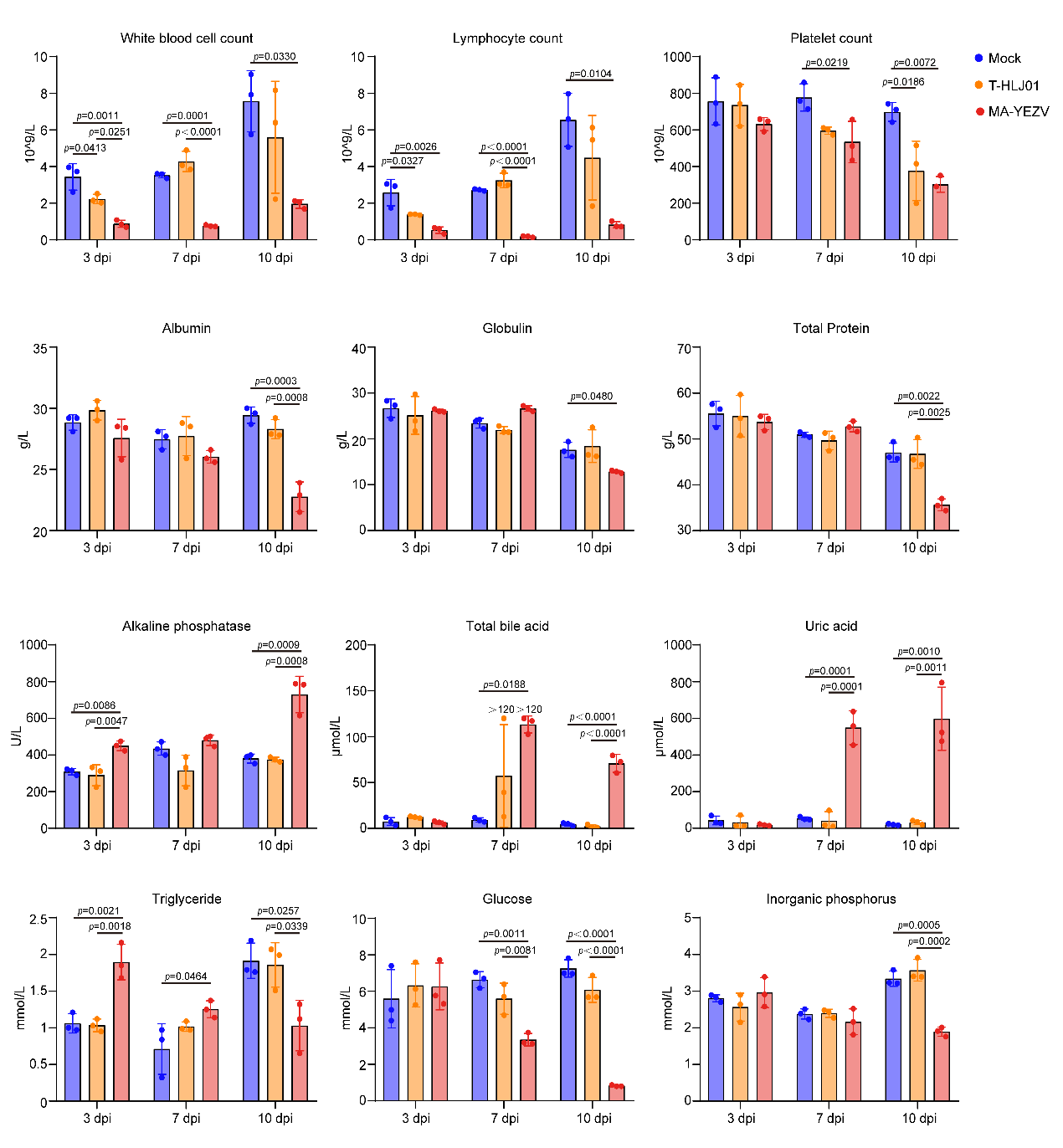
**Supplementary Fig. 7 Hematological and biochemical analysis of C57BL/6J mice infected with T-HLJ01 and MA-YEZV**

Blood was collected form male C57BL/6J mice at 3, 7, and 10 days post-infection (dpi) following intraperitoneal inoculation with MA-YEZV or the T-HLJ01 strain. Key parameters are shown: white blood cell count, lymphocyte count, platelet count, albumin, globulin, total protein, alkaline phosphatase, total bile acid, uric acid, triglycerides, glucose, and inorganic phosphorus. Data are presented as mean ± standard deviation (n = 3 biological replicates). Statistical significance was determined by one-way ANOVA (two-sided) followed by Tukey’s multiple-comparison test to adjust for multiple testing. Exact *p* values are provided in the figure legends. Source data are provided as a Source Data file.

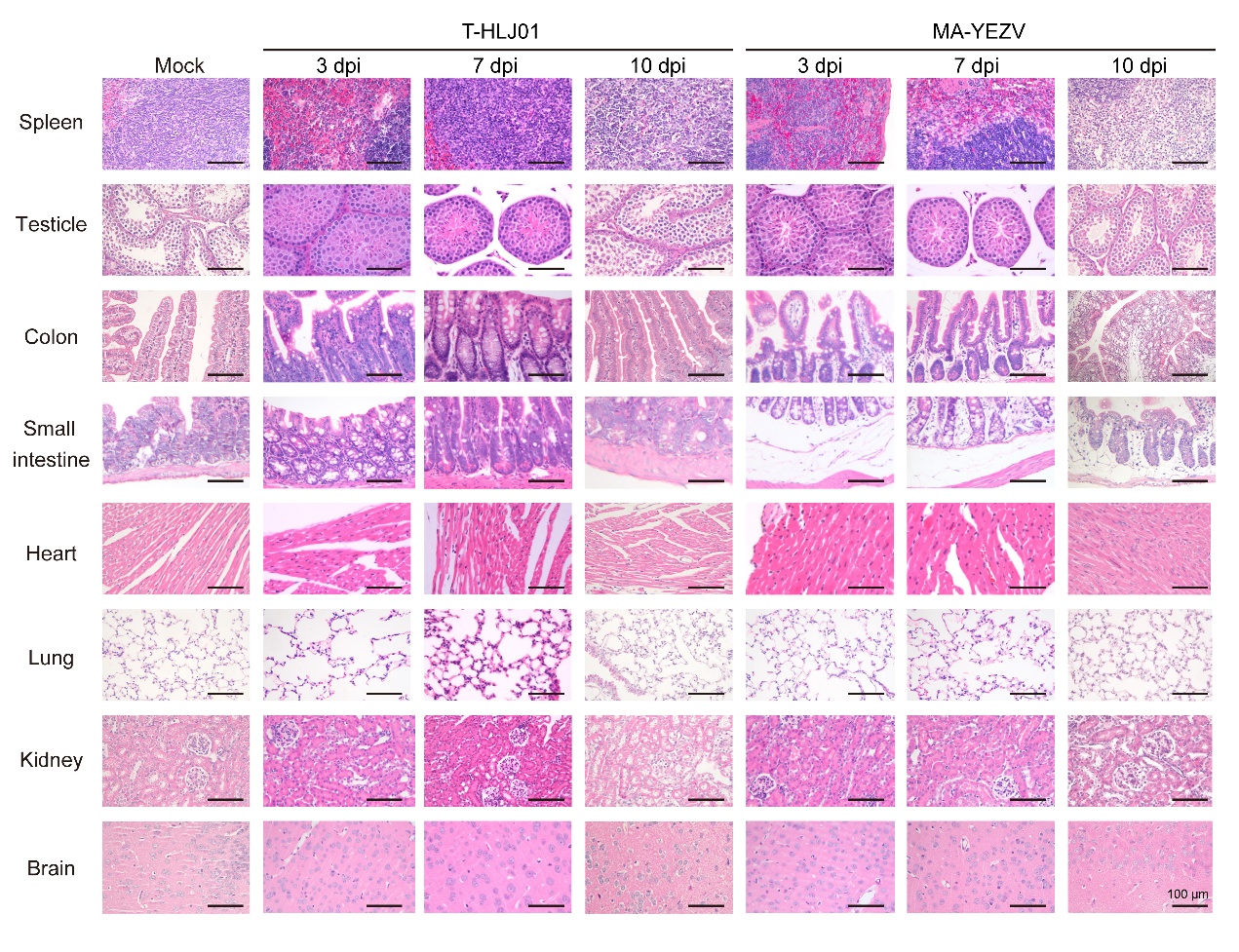

**Supplementary Fig. 8 Comparative tissue pathology of C57BL/6J mice infected with T-HLJ01 and MA-YEZV.**

Representative hematoxylin and eosin (H&E)-stained sections of multiple tissues harvested at 3, 7, and 10 days post-infection (dpi) from mice intraperitoneally inoculated with MA-YEZV (1 TCID_50_) or T-HLJ01 (10^4^ TCID_50_). Tissues shown include the spleen, testicle, colon, small intestine, heart, lung, kidney, and brain. Images illustrate the broad tissue damage and inflammatory infiltration induced by MA-YEZV, in contrast to the minimal pathology observed with the parental T-HLJ01 strain. Scale bar, 100 µm. Experiments were performed on three independent biological replicates, and representative images are shown.

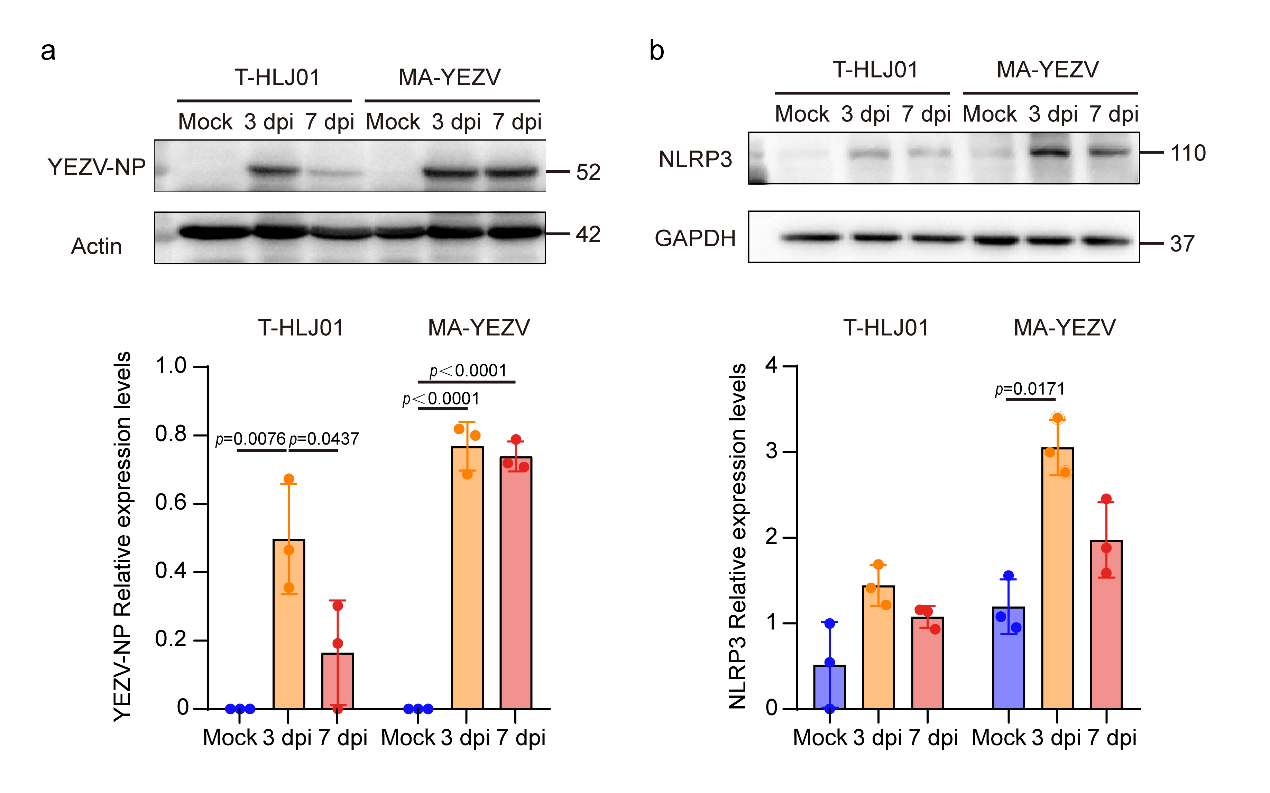
**Supplementary Fig. 9 MA-YEZV infection induces sustained hepatic inflammasome activation and viral protein expression.**

(a) Representative immunoblots and quantitative analysis of the viral nucleoprotein (NP) in the liver lysates from C57BL/6J mice infected with 1 TCID_50_ of MA-YEZV or 10^4^ TCID₅₀ of the T-HLJ01 strain at 3 and 7 days post-infection (dpi). Actin served as the loading control.

(b) Representative immunoblots and quantitative analysis of the inflammasome sensor NLRP3 in the liver lysates from C57BL/6J mice infected with 1 TCID_50_ of MA-YEZV or 10^4^ TCID₅₀ of the T-HLJ01 strain at 3 and 7 days post-infection (dpi). GAPDH served as the loading control.

Data are presented as mean ± standard deviation (n = 3 biological replicates). Statistical significance was determined by one-way ANOVA (two-sided) followed by Tukey’s multiple-comparison test to adjust for multiple testing. Exact *p* values are provided in the figure legends.

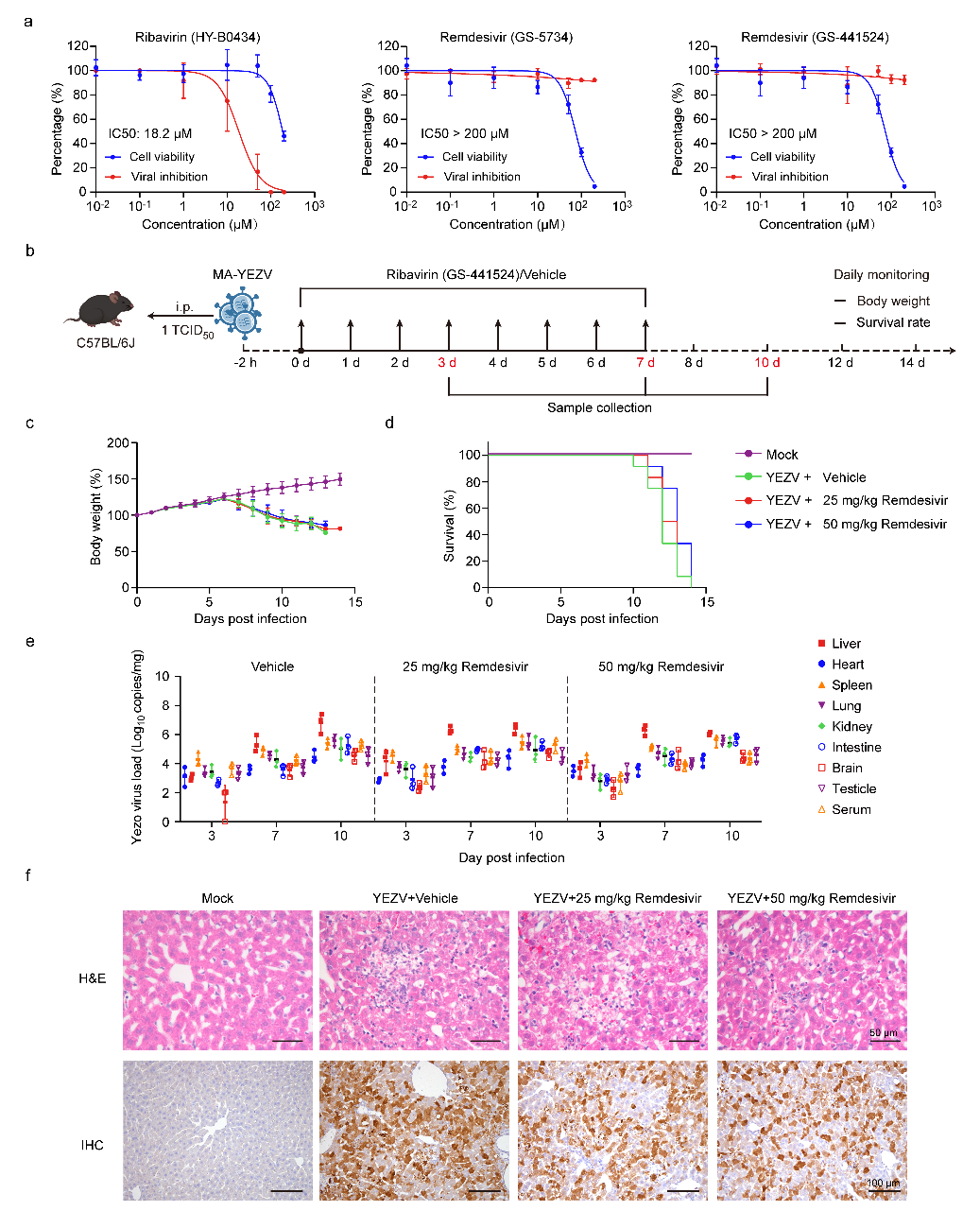
**Supplementary Fig. 10 The remdesivir metabolite GS-441524 lacks antiviral efficacy against MA-YEZV**

(a) *In vitro* efficacy and cytotoxicity. Vero cells infected with MA-YEZV (MOI 0.01) were treated with the indicated concentrations of ribavirin (HY-B0434), remdesivir (GS-5734), or the remdesivir metabolite (GS-441524). Supernatant viral titers at 5 dpi and compound cytotoxicity (CCK-8 assay) are shown (n=3 biological replicates).

(b) Schematic of the *in vivo* treatment protocol. Mice infected with MA-YEZV (1 TCID₅₀, i.p.) were treated with GS-441524 (25 or 50 mg/kg) or vehicle daily for 8 days.

(c, d) Body weight change (b) and survival (c) of infected mice treated with GS-441524 or vehicle (n=12 independent mice, n=6 independent mice).

(e) Viral loads in tissues and serum of infected mice treated with GS-441524 or vehicle (n=3 biological replicates) measured by RT-qPCR at 3, 7, and 10 days post-infection.

(f) Histopathology of liver at 10 dpi (H&E) and viral antigen detection (IHC). Scale bars, 50 µm (H&E), 100 µm (IHC). Experiments were performed on three independent biological replicates, and representative images are shown.

Data in (a, c, and e) are mean ± standard deviation. Source data are provided as a Source Data file.

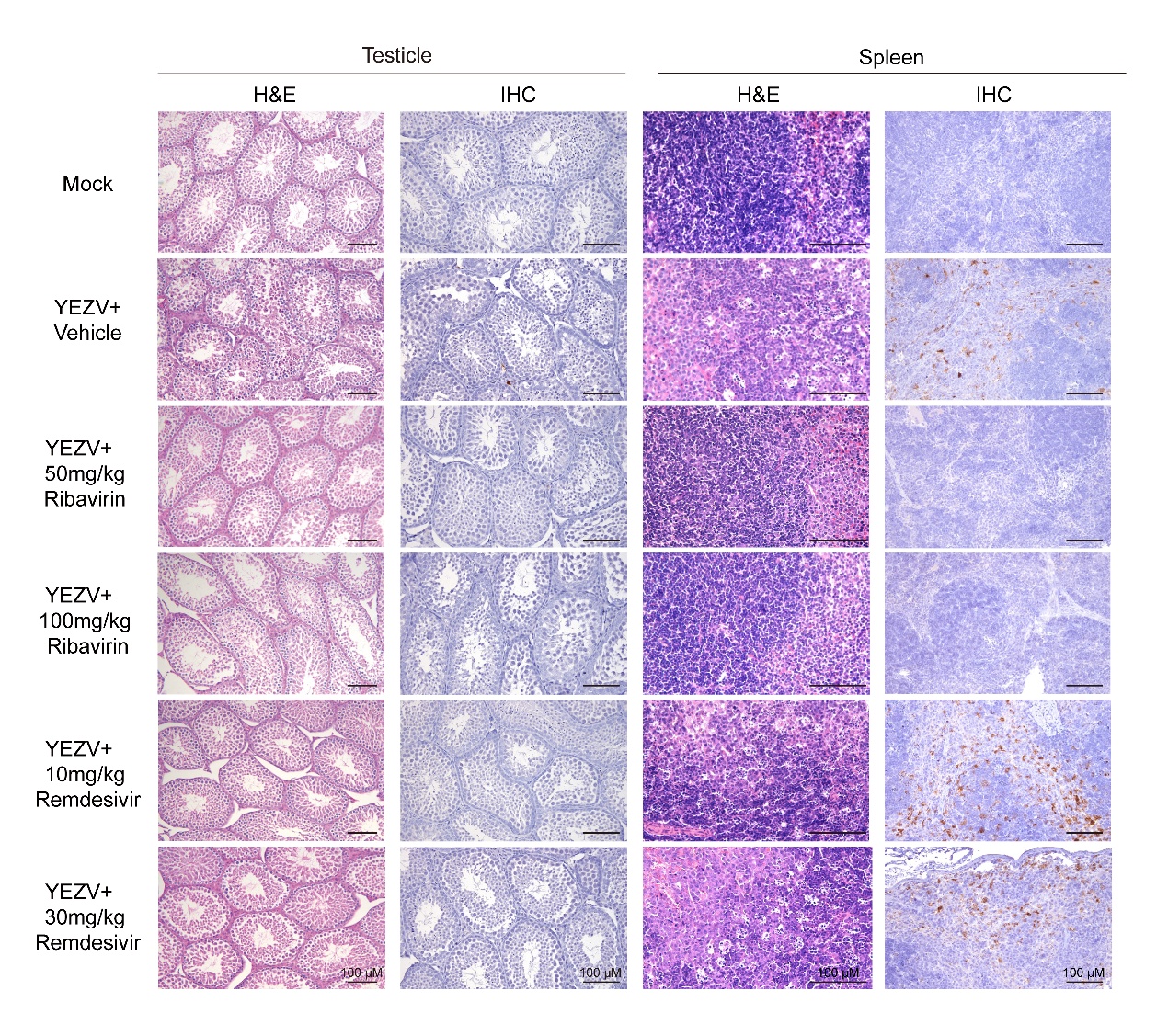
 **Supplementary Fig. 11 Histopathological analysis of spleen and testes in MA-YEZV-infected C57BL/6J mice treated with antiviral drugs.**

Three-week-old male C57BL/6J mice were intraperitoneally inoculated with 1 TCID_50_ of MA-YEZV. Two hours post-infection, mice were treated with either ribavirin (50 or 100 mg/kg) or remdesivir (10 or 30 mg/kg), while vehicle-treated mice served as controls. Antiviral treatments were administered once daily for a total of 8 consecutive days. Ten days post-infection (dpi), tissues were collected for pathological analysis. H&E staining and immunohistochemical (IHC) staining of spleen and testes from C57BL/6J mice at 10 dpi following treatment with ribavirin, remdesivir, or vehicle after MA-YEZV infection. Scale bars: 100 μm (H&E), 50 μm (IHC). Experiments were performed on three independent biological replicates, and representative images are shown.

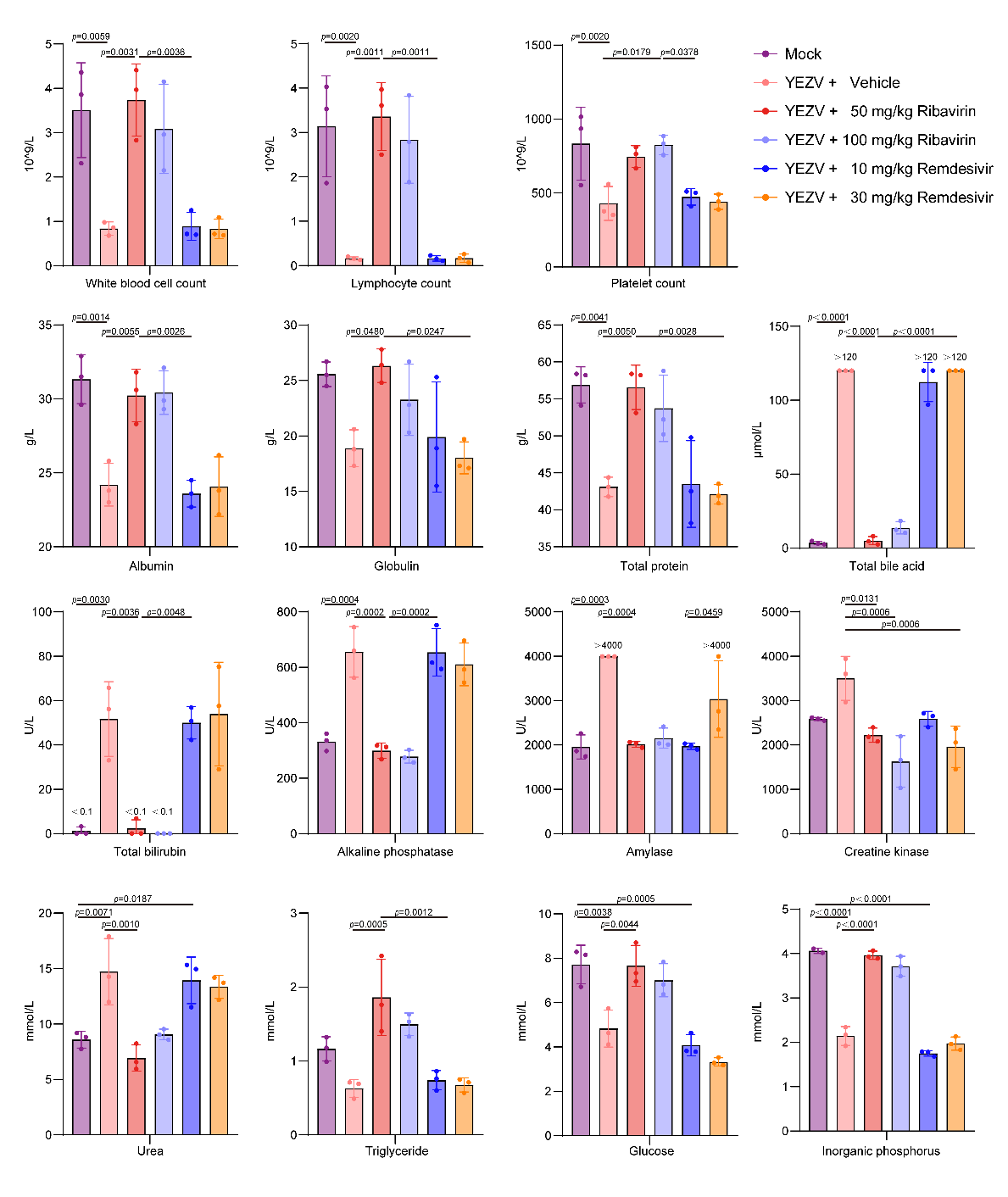
**Supplementary Fig. 12 Hematological and biochemical analysis of MA-YEZV-infected C57BL/6J mice following antiviral treatment**

Three-week-old male C57BL/6J mice were intraperitoneally inoculated with 1 TCID_50_ of MA-YEZV. Two hours post-infection, mice were treated with either ribavirin (50 or 100 mg/kg) or remdesivir (10 or 30 mg/kg), while vehicle-treated mice served as controls. Antiviral treatments were administered once daily for a total of 8 consecutive days. Ten days post-infection, blood samples were collected for hematological and biochemical analyses. The detected parameters include white blood cell count, lymphocyte count, platelet count, albumin, globulin, total protein, total bile acid, total bilirubin, alkaline phosphatase, amylase, creatine kinase, urea, triglyceride, glucose and inorganic phosphorus. Data are presented as mean ± standard deviation (n = 3 biological replicates). Statistical significance was determined by one-way ANOVA (two-sided) followed by Tukey’s multiple-comparison test to adjust for multiple testing. Exact *p* values are provided in the figure legends. Source data are provided as a Source Data file.

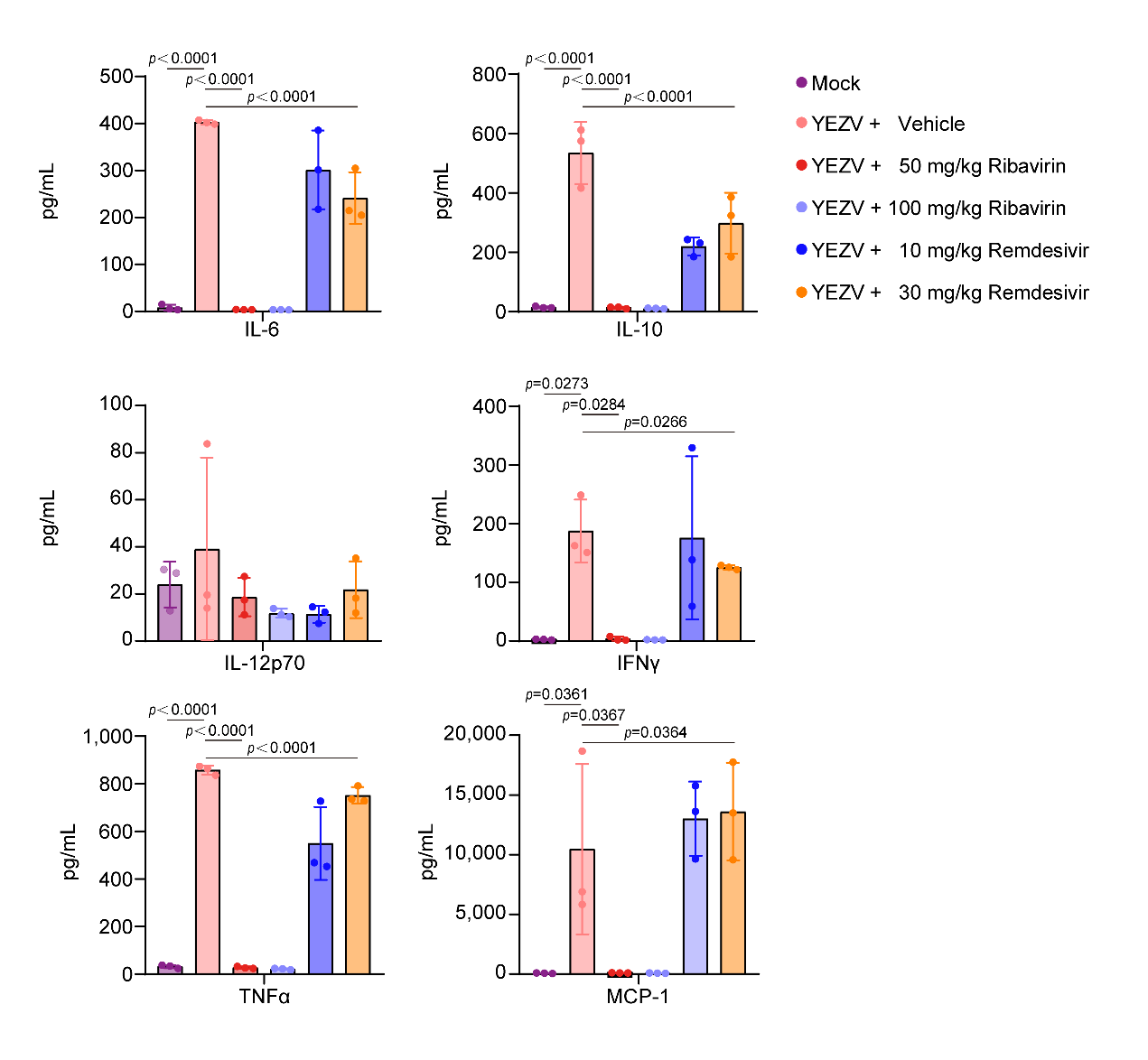
**Supplementary Fig. 13 Hematological and biochemical analysis of MA-YEZV-infected C57BL/6J mice following antiviral treatment**

Three-week-old male C57BL/6J mice were intraperitoneally inoculated with 1 TCID_50_ of MA-YEZV. Two hours post-infection, mice were treated with either ribavirin (50 or 100 mg/kg) or remdesivir (10 or 30 mg/kg), while vehicle-treated mice served as controls. Antiviral treatments were administered once daily for a total of 8 consecutive days. Ten days post-infection, blood samples were collected for cytokine analysis. Cytokines in serum were measured using the CBA method. The detected parameters included IL-6, IL-10, IL-12p70, IFN-γ, MCP-1, and TNF-α. Data are presented as mean ± standard deviation (n = 3 biological replicates). Statistical significance was determined by one-way ANOVA (two-sided) followed by Tukey’s multiple-comparison test to adjust for multiple testing. Exact *p* values are provided in the figure legends. Source data are provided as a Source Data file.

**Supplementary Table 1** Nucleotide mutation sites analysis of MA-YEZV.

| **Viral genome** | **Mutation** | **Synonymous mutation** | **Nonsynonymous mutation** |
| --- | --- | --- | --- |
| L-5’ UTR* | 10 | — | — |
| L-CDS† | 167 | 153 | 14 |
| L-3’ UTR | 1 | — | — |
| M-5’ UTR | 1 | — | — |
| M-CDS | 68 | 57 | 11 |
| M-3’ UTR | 1 | — | — |
| S-5’ UTR | 0 | — | — |
| S-CDS | 107 | 102 | 5 |
| S-3’ UTR | 12 | — | — |

* UTR, untranslated region; † CDS, coding sequence.

**Supplementary Table 2** Amino acid mutation sites analysis of MA-YEZV.

| **Segment** | **Mutation sites** | | **Position (aa)** | **Mutation Site Comparison** | |
| --- | --- | --- | --- | --- | --- |
|  | **T-HLJ01** | **MA**-**YEZV** |  | **Human-derived (n=13)** | **Tick-derived (n=29)** |
| L | E | G | 83 | 7.70% (1/13) | 13.79% (4/29) |
|  | N | K | 152 | Not reported | Not reported |
|  | A | S | 163 | Not reported | Not reported |
|  | D | E | 495 | 100.00% (13/13) | 89.66% (26/29) |
|  | R | K | 520 | 92.31% (12/13) | 96.55% (28/29) |
|  | V | I | 965 | 61.54% (8/13) | 48.28% (14/29) |
|  | K | R | 1188 | Not reported | 3.45% (1/29) |
|  | K | R | 2088 | 92.31% (12/13) | 72.41% (21/29) |
|  | L | M | 2095 | 100.00% (13/13) | 100.00% (29/29) |
|  | R | K | 2110 | 69.23% (9/13) | 44.83% (13/29) |
|  | T | S | 2407 | Not reported | 3.45% (1/29) |
|  | V | I | 3518 | Not reported | 3.45% (1/29) |
|  | V | I | 3899 | 7.69% (1/13) | 6.90% (2/29) |
| M | C | Y | 6 | 7.69% (1/13) | 20.69% (6/29) |
|  | E | G | 22 | Not reported | Not reported |
|  | E | K | 56 | Not reported | Not reported |
|  | D | N | 190 | Not reported | 3.45% (1/29) |
|  | S | F | 244 | Not reported | Not reported |
|  | T | A | 728 | 7.69% (1/13) | 24.14% (7/29) |
|  | L | V | 729 | 69.23% (9/13) | 44.83% (13/29) |
|  | C | S | 873 | 100.00% (13/13) | 96.55% (28/29) |
|  | Q | R | 1086 | Not reported | Not reported |
|  | I | L | 1261 | Not reported | Not reported |
|  | R | K | 1332 | Not reported | 3.45% (1/29) |
| S | C | Y | 46 | Not reported | Not reported |
|  | K | M | 301 | Not reported | Not reported |
|  | T | A | 498 | 46.15% (6/13) | 13.79% (4/29) |
|  | K | T | 501 | 38.46% (5/13) | 13.79% (4/29) |
|  | P | H | 502 | 38.46% (5/13) | 20.69% (6/29) |

**Supplementary Table 3** Blood biochemical analysis of C57BL/6J mice infected with MA-YEZV at 3 dpi.

| **Test items** | **Mock** | **T-HLJ01** | **MA-YEZV** | **Reference value** |
| --- | --- | --- | --- | --- |
| Albumin (g/L) | 28.83±0.52 | 29.83±0.65 | 27.57±1.25 | 25.0-48.0 |
| Globulin (g/L) | 26.67±1.65 | 25.10±3.37 | 26.13±0.26 | — |
| Total protein (g/L) | 55.53±2.19 | 54.97±3.69 | 53.70±1.36 | 36.0-72.0 |
| Total bilirubin (μmol/L) | <0.50±0.57 | 0.33±0.09 | 1.46±1.14 | 0.0-15.0 |
| Glutamyl transferase (U/L) | <2±0.00 | <2±0.00 | <2±0.00 | — |
| Alanine aminotransferase (U/L) | 205.33±41.43 | 172.33±8.96 | 356.00±5.10 | 59-247 |
| Aspartate aminotransferase (U/L) | 78.00±15.30 | 69.33±1.89 | 248.67±35.83 | 28-132 |
| Alkaline phosphatase (U/L) | 308.67±14.20 | 289.67±46.45 | 450.00±21.28 | 62-209 |
| Total bile acid (μmol/L) | 7.51±3.49 | 12.20±0.85 | 6.32±1.40 | — |
| Amylase (U/L) | 2005.00±36.34 | 1863.00±222.76 | 1577.67±18.37 | 1691-3615 |
| Lipase (U/L) | 61.33±5.25 | 35.23±8.95 | 45.67±5.79 | — |
| Lactate dehydrogenase (U/L) | 976.33±136.71 | 984.67±44.13 | 1030.00±28.25 | 1105-3993 |
| Creatine kinase (U/L) | 2402.33±501.38 | 2108.33±55.70 | 2752.00±826.60 | 68-1070 |
| Creatinine (μmol/L) | 26.30±5.36 | 17.03±4.46 | 17.53±2.98 | 12.0-71.0 |
| Uric acid (μmol/L) | 43.16±18.90 | 32.97±27.90 | 18.32±3.63 | 101.0-321.0 |
| Urea (mmol/L) | 7.91±0.39 | 8.07±0.54 | 8.69±0.28 | 4.0-11.8 |
| Glucose (mmol/L) | 5.59±1.30 | 6.32±0.97 | 6.26±1.05 | 5.0-10.7 |
| Total cholesterol (mmol/L) | 2.50±0.23 | 2.24±0.34 | 2.39±0.10 | 0.93-4.04 |
| Triglyceride (mmol/L) | 1.06±0.11 | 1.03±0.07 | 1.90±0.20 | 0.62-1.63 |
| Total carbon dioxide (mmol/L) | 20.93±0.63 | 16.90±2.53 | 20.77±1.18 | — |
| Calcium (mmol/L) | 2.51±0.03 | 1.46±0.76 | 2.51±0.10 | 1.48-2.35 |
| Inorganic phosphorus (mmol/L) | 2.81±0.08 | 2.56±0.31 | 2.96±0.33 | 1.97-3.26 |

**Supplementary Table 4** Hematological analysis of C57BL/6J mice infected with MA-YEZV at 3 dpi.

| **Test items** | **Mock** | **T-HLJ01** | **MA-YEZV** | **Reference value** |
| --- | --- | --- | --- | --- |
| White blood cell count (10^9/L) | 3.43±0.59 | 2.23±0.21 | 0.88±0.16 | 3.61-13.00 |
| Neutrophil count (10^9/L) | 0.75±0.06 | 0.72±0.12 | 0.32±0.06 | 0.10-2.00 |
| Lymphocyte count (10^9/L) | 2.58±0.59 | 1.38±0.02 | 0.53±0.14 | 1.27-8.44 |
| Monocyte count (10^9/L) | 0.06±0.01 | 0.13±0.08 | 0.01±0.01 | 0.00-0.29 |
| Eosinophil count (10^9/L) | 0.03±0.03 | 0.00±0.00 | 0.02±0.01 | 0.00-0.17 |
| Basophil count (10^9/L) | 0.01±0.00 | 0.00±0.00 | 0.00±0.00 | 0.00-0.20 |
| Neutrophil percentage (%) | 22.20±2.95 | 31.90±2.57 | 36.07±7.66 | 1.2-30.0 |
| Lymphocyte percentage (%) | 73.93±5.15 | 62.17±5.48 | 58.97±9.84 | 70.0-96.0 |
| Monocyte percentage (%) | 2.03±0.91 | 5.23±3.28 | 2.13±1.33 | 0.0-10.0 |
| Eosinophil percentage (%) | 1.33±1.27 | 0.40±0.22 | 2.17±0.47 | 0.0-10.0 |
| Basophil percentage (%) | 0.50±0.08 | 0.30±0.08 | 0.67±0.53 | 0.0-5.0 |
| Red blood cell count (10^12/L) | 7.93±0.30 | 8.53±0.46 | 8.15±0.21 | 6.00-12.50 |
| Hemoglobin concentration (g/L) | 130.00±7.35 | 127.33±8.96 | 133.67±4.64 | 100-190 |
| Hematocrit (%) | 36.70±1.61 | 38.77±2.04 | 36.60±0.83 | 40.0-48.0 |
| Mean corpuscular volume (fL) | 46.27±0.78 | 45.50±0.22 | 44.90±0.29 | 41.0-63.0 |
| Mean corpuscular hemoglobin (pg) | 16.37±0.34 | 14.97±1.75 | 16.37±0.17 | 13.0-19.0 |
| Mean corpuscular hemoglobin concentration (g/L) | 353.33±6.18 | 329.67±38.00 | 364.67±6.02 | 290-351 |
| Coefficient of variation of red cell distribution width (%) | 18.60±0.99 | 18.27±0.12 | 17.27±0.48 | 10.0-20.0 |
| Red cell distribution width standard deviation (fL) | 33.83±1.65 | 32.23±0.45 | 30.17±1.11 | 0.1-99.9 |
| Platelet count (10^9/L) | 756.00±104.75 | 735.67±91.96 | 632.00±29.34 | 540-154 |
| Mean platelet volume (fL) | 9.80±0.75 | 11.07±0.05 | 9.80±0.33 | 3.8-14.1 |
| Platelet distribution width (fL) | 7.03±0.94 | 7.13±0.80 | 5.53±1.18 | 0.1-30.0 |
| Platelet hematocrit (%) | 0.74±0.11 | 0.81±0.10 | 0.72±0.11 | 0.10-9.99 |

**Supplementary Table 5** Blood biochemical analysis of C57BL/6J mice infected with MA-YEZV 7 dpi.

| **Test items** | **Mock** | **T-HLJ01** | **MA-YEZV** | **Reference value** |
| --- | --- | --- | --- | --- |
| Albumin (g/L) | 27.47±0.63 | 27.73±1.30 | 26.03±0.42 | 25.0-48.0 |
| Globulin (g/L) | 23.40±0.86 | 21.93±0.61 | 26.60±0.49 | — |
| Total protein (g/L) | 50.90±0.45 | 49.67±1.65 | 52.70±0.94 | 36.0-72.0 |
| Total bilirubin (μmol/L) | <0.1±0.00 | 2.17±0.69 | 17.97±6.41 | 0.0-15.0 |
| Glutamyl transferase (U/L) | <2±0.00 | <2±0.00 | <2±0.00 | — |
| Alanine aminotransferase (U/L) | 159.33±8.73 | 214.33±25.25 | >650±0.00 | 59-247 |
| Aspartate aminotransferase (U/L) | 66.00±5.66 | 85.67±13.47 | >650±0.00 | 28-132 |
| Alkaline phosphatase (U/L) | 435.00±29.43 | 315.00±67.05 | 479.67±23.61 | 62-209 |
| Total bile acid (μmol/L) | 9.26±1.95 | 25.99±13.13 | 109.96±7.21 | — |
| Amylase (U/L) | 898.33±122.92 | 686.67±170.36 | 1077.33±56.75 | 1691-3615 |
| Lipase (U/L) | 63.67±14.34 | 49.33±6.34 | 53.33±3.40 | — |
| Lactate dehydrogenase (U/L) | 603.33±43.05 | 685.67±72.69 | >4000±0.00 | 1105-3993 |
| Creatine kinase (U/L) | 2185.33±345.75 | 2680.33±453.90 | 2725.67±472.97 | 68-1070 |
| Creatinine (μmol/L) | 19.43±1.83 | 13.00±2.00 | 12.20±0.00 | 12.0-71.0 |
| Uric acid (μmol/L) | 53.77±7.21 | 57.51±40.57 | 549.31±75.20 | 101.0-321.0 |
| Urea (mmol/L) | 8.73±0.97 | 7.80±0.72 | 11.72±1.53 | 4.0-11.8 |
| Glucose (mmol/L) | 6.63±0.36 | 5.59±0.69 | 3.35±0.28 | 5.0-10.7 |
| Total cholesterol (mmol/L) | 2.26±0.14 | 1.62±0.12 | 3.22±0.11 | 0.93-4.04 |
| Triglyceride (mmol/L) | 0.71±0.28 | 1.02±0.05 | 1.25±0.09 | 0.62-1.63 |
| Total carbon dioxide (mmol/L) | 15.60±0.14 | 15.10±0.28 | 19.53±1.39 | — |
| Calcium (mmol/L) | 2.93±0.01 | 2.94±0.03 | 2.19±0.02 | 1.48-2.35 |
| Inorganic phosphorus (mmol/L) | 2.38±0.11 | 2.39±0.09 | 2.16±0.29 | 1.97-3.26 |

**Supplementary Table 6** Hematological analysis of C57BL/6J mice infected with MA-YEZV 7 dpi.

| **Test items** | **Mock** | **T-HLJ01** | **MA-YEZV** | **Reference value** |
| --- | --- | --- | --- | --- |
| White blood cell count (10^9/L) | 3.52±0.11 | 4.27±0.45 | 0.76±0.04 | 3.61-13.00 |
| Neutrophil count (10^9/L) | 0.73±0.09 | 0.95±0.15 | 0.50±0.03 | 0.10-2.00 |
| Lymphocyte count (10^9/L) | 2.72±0.04 | 3.24±0.32 | 0.18±0.04 | 1.27-8.44 |
| Monocyte count (10^9/L) | 0.05±0.01 | 0.05±0.00 | 0.05±0.00 | 0.00-0.29 |
| Eosinophil count (10^9/L) | 0.01±0.01 | 0.01±0.01 | 0.02±0.01 | 0.00-0.17 |
| Basophil count (10^9/L) | 0.00±0.00 | 0.02±0.01 | 0.01±0.01 | 0.00-0.20 |
| Neutrophil percentage (%) | 20.53±1.92 | 22.03±1.44 | 65.43±4.17 | 1.2-30.0 |
| Lymphocyte percentage (%) | 77.37±1.43 | 75.93±1.30 | 22.63 ± 4.20 | 70.0-96.0 |
| Monocyte percentage (%) | 1.40±0.28 | 1.27±0.29 | 7.40±1.47 | 0.0-10.0 |
| Eosinophil percentage (%) | 0.47±0.26 | 0.30±0.22 | 2.77±1.54 | 0.0-10.0 |
| Basophil percentage (%) | 0.23±0.12 | 0.50±0.22 | 1.77±1.07 | 0.0-5.0 |
| Red blood cell count (10^12/L) | 7.81±0.90 | 7.99±0.27 | 8.44±0.17 | 6.00-12.50 |
| Hemoglobin concentration (g/L) | 128.33±14.82 | 130.33±3.30 | 136.33±4.19 | 100-190 |
| Hematocrit (%) | 35.53±4.46 | 37.07±0.66 | 37.97±1.39 | 40.0-48.0 |
| Mean corpuscular volume (fL) | 45.43±1.38 | 46.43±0.71 | 44.93±0.75 | 41.0-63.0 |
| Mean corpuscular hemoglobin (pg) | 16.43±0.12 | 16.30±0.14 | 16.13±0.21 | 13.0-19.0 |
| Mean corpuscular hemoglobin concentration (g/L) | 361.67±7.93 | 351.33±2.49 | 359.00±2.45 | 290-351 |
| Coefficient of variation of red cell distribution width (%) | 19.37±2.23 | 18.10±0.78 | 16.83±0.60 | 10.0-20.0 |
| Red cell distribution width standard deviation (fL) | 34.80±5.34 | 32.97±2.04 | 29.47±1.41 | 0.1-99.9 |
| Platelet count (10^9/L) | 843.67±153.32 | 595.33±16.21 | 534.33±92.40 | 540-154 |
| Mean platelet volume (fL) | 9.93±0.80 | 8.70±0.08 | 11.50±0.65 | 3.8-14.1 |
| Platelet distribution width (fL) | 6.53±0.94 | 5.37±0.24 | 7.17±0.05 | 0.1-30.0 |
| Platelet hematocrit (%) | 0.85±0.20 | 0.68±0.04 | 0.61±0.10 | 0.10-9.99 |

**Supplementary Table 7** Blood biochemical analysis of C57BL/6J mice infected with MA-YEZV 10 dpi.

| **Test items** | **Mock** | **T-HLJ01** | **MA-YEZV** | **Reference value** |
| --- | --- | --- | --- | --- |
| Albumin (g/L) | 29.43±0.54 | 28.30±0.64 | 22.77±0.98 | 25.0-48.0 |
| Globulin (g/L) | 17.60±1.36 | 18.40±2.90 | 12.83±0.25 | — |
| Total protein (g/L) | 47.03±1.68 | 46.73±2.56 | 35.63±1.07 | 36.0-72.0 |
| Total bilirubin (μmol/L) | <5.43±5.21 | <6.87±9.57 | <3.23±4.43 | 0.0-15.0 |
| Glutamyl transferase (U/L) | <2±0.00 | <2±0.00 | <5.33±2.87 | — |
| Alanine aminotransferase (U/L) | 168.00±4.32 | 210.33±8.65 | >650±0.00 | 59-247 |
| Aspartate aminotransferase (U/L) | 59.00±7.87 | 83.00±11.43 | >650±0.00 | 28-132 |
| Alkaline phosphatase (U/L) | 380.33±19.75 | 375.00±10.23 | 729.33±80.87 | 62-209 |
| Total bile acid (μmol/L) | 4.50±1.36 | <4.13±1.00 | 70.90±7.94 | — |
| Amylase (U/L) | 2112.00±203.54 | 1700.67±119.21 | 2216.33±762.98 | 1691-3615 |
| Lipase (U/L) | 54.67±7.41 | 56.33±2.36 | 51.00±1.63 | — |
| Lactate dehydrogenase (U/L) | 735.33±114.11 | 1086.33±340.05 | 3374.00±607.04 | 1105-3993 |
| Creatine kinase (U/L) | 2478.33±280.52 | 2768.33±749.51 | 2618.00±240.94 | 68-1070 |
| Creatinine (μmol/L) | <11.60±1.99 | <10.67±0.94 | <10.0±0.00 | 12.0-71.0 |
| Uric acid (μmol/L) | 20.71±3.70 | 32.12±9.08 | 597.76±140.86 | 101.0-321.0 |
| Urea (mmol/L) | 10.55±0.16 | 10.32±0.81 | 9.92±0.71 | 4.0-11.8 |
| Glucose (mmol/L) | 7.25±0.39 | 6.07±0.56 | <0.82±0.03 | 5.0-10.7 |
| Total cholesterol (mmol/L) | 2.09±0.06 | 2.70±0.22 | 2.12±0.13 | 0.93-4.04 |
| Triglyceride (mmol/L) | 1.91±0.20 | 1.86±0.25 | 1.03±0.28 | 0.62-1.63 |
| Total carbon dioxide (mmol/L) | 20.27±0.94 | 20.60±1.12 | 20.00±1.14 | — |
| Calcium (mmol/L) | 2.45±0.09 | 2.36±0.07 | 2.21±0.18 | 1.48-2.35 |
| Inorganic phosphorus (mmol/L) | 3.34±0.17 | 3.57±0.24 | 3.89±0.10 | 1.97-3.26 |

**Supplementary Table 8** Hematological analysis of C57BL/6J mice infected with MA-YEZV 10 dpi.

| **Test items** | **Mock** | **T-HLJ01** | **MA-YEZV** | **Reference value** |
| --- | --- | --- | --- | --- |
| White blood cell count (10^9/L) | 7.57±1.36 | 5.60±2.49 | 1.96±0.19 | 3.61-13.00 |
| Neutrophil count (10^9/L) | 0.92±0.21 | 0.74±0.35 | 0.62±0.12 | 0.10-2.00 |
| Lymphocyte count (10^9/L) | 6.54±1.18 | 4.48±1.88 | 0.83±0.14 | 1.27-8.44 |
| Monocyte count (10^9/L) | 0.08±0.06 | 0.38±0.34 | 0.39±0.21 | 0.00-0.29 |
| Eosinophil count (10^9/L) | 0.01±0.01 | 0.00±0.00 | 0.09±0.02 | 0.00-0.17 |
| Basophil count (10^9/L) | 0.02±0.01 | 0.01±0.01 | 0.02±0.01 | 0.00-0.20 |
| Neutrophil percentage (%) | 12.10±1.78 | 13.10±1.27 | 31.50±3.63 | 1.2-30.0 |
| Lymphocyte percentage (%) | 86.47±2.31 | 80.93±4.33 | 41.93±5.57 | 70.0-96.0 |
| Monocyte percentage (%) | 0.93±0.69 | 5.77±3.35 | 19.87±9.45 | 0.0-10.0 |
| Eosinophil percentage (%) | 0.10±0.08 | 0.07±0.05 | 5.17±1.92 | 0.0-10.0 |
| Basophil percentage (%) | 0.40±0.22 | 0.17±0.05 | 1.53±0.54 | 0.0-5.0 |
| Red blood cell count (10^12/L) | 7.77±0.01 | 5.80±1.03 | 7.59±0.10 | 6.00-12.50 |
| Hemoglobin concentration (g/L) | 122.33±10.21 | 95.33±19.15 | 125.00±1.63 | 100-190 |
| Hematocrit (%) | 37.67±0.49 | 28.37±5.29 | 35.93±0.26 | 40.0-48.0 |
| Mean corpuscular volume (fL) | 48.50±0.70 | 48.83±0.85 | 47.37±0.31 | 41.0-63.0 |
| Mean corpuscular hemoglobin (pg) | 15.70±1.36 | 16.37±0.45 | 16.50±0 | 13.0-19.0 |
| Mean corpuscular hemoglobin concentration (g/L) | 323.67±23.84 | 335.00±7.12 | 348.00±2.83 | 290-351 |
| Coefficient of variation of red cell distribution width (%) | 18.10±1.50 | 18.00±1.20 | 17.97±0.97 | 10.0-20.0 |
| Red cell distribution width standard deviation (fL) | 35.07±2.35 | 35.63±1.77 | 33.90±1.69 | 0.1-99.9 |
| Platelet count (10^9/L) | 698.33±41.80 | 376.33±131.87 | 303.00±35.33 | 540-154 |
| Mean platelet volume (fL) | 8.40±0.67 | 8.77±0.37 | 12.13±0.25 | 3.8-14.1 |
| Platelet distribution width (fL) | 5.97±0.62 | 8.47±0.62 | 9.10±1.57 | 0.1-30.0 |
| Platelet hematocrit (%) | 0.59±0.08 | 0.33±0.12 | 0.37±0.05 | 0.10-9.99 |

**Supplementary Table 9** Blood biochemical analysis of C57BL/6J mice infected with MA-YEZV following antiviral drug treatment.

| **Test items** | **Mock** | **YEZV** | **YEZV+50 mg/kg Ribavirin** | **YEZV+100 mg/kg Ribavirin** | **YEZV+10 mg/kg Remdesivir** | **YEZV+30 mg/kg Remdesivir** | **Reference value** |
| --- | --- | --- | --- | --- | --- | --- | --- |
| Albumin (g/L) | 31.33±1.65 | 24.87±1.44 | 30.23±1.26 | 30.43±1.19 | 23.60±0.90 | 24.07±2.01 | 25.0-48.0 |
| Globulin (g/L) | 25.57±1.10 | 18.90±1.76 | 26.33±1.72 | 23.27±3.22 | 19.90±4.99 | 18.03±1.15 | — |
| Total protein (g/L) | 56.90±2.32 | 43.10±1.30 | 56.57±0.14 | 53.73±4.54 | 43.50±5.83 | 42.10±1.31 | 36.0-72.0 |
| Total bilirubin (μmol/L) | <1.17±1.86 | 51.70±14.22 | <0.07±0.05 | <0.10±0.00 | 50.00±7.36 | 53.97±23.31 | 0.0-15.0 |
| Glutamyl transferase (U/L) | <2.00±0.00 | ＜2.00±0.00 | <2.00±0.00 | <2.00±0.00 | ＜2.00±0.00 | ＜2.00±0.00 | — |
| Alanine aminotransferase (U/L) | 209.00±33.17 | ＞650.00±0.00 | 154.67±28.06 | 163.00±51.86 | ＞650.00±0.00 | ＞650.00±0.00 | 59-247 |
| Aspartate aminotransferase (U/L) | 85.33±8.50 | ＞650.00±0.00 | 60.00±10.01 | 71.00±14.42 | ＞650.00±0.00 | ＞650.00±0.00 | 28-132 |
| Alkaline phosphatase (U/L) | 331.67±32.29 | 655.67±92.26 | 299.67±25.17 | 278.00±23.88 | 654.33±84.27 | 611.00±75.68 | 62-209 |
| Total bile acid (μmol/L) | 3.50±1.15 | ＞120.00±0.00 | 5.71±2.79 | 13.71±4.25 | ＞112.33±13.27 | ＞120.00±0.00 | — |
| Amylase (U/L) | 1956.00±267.25 | ＞4000.00±0.00 | 2029.00±56.19 | 2154.00±220.92 | 1972.00±68.29 | ＞3035.00±873.56 | 1691-3615 |
| Lipase (U/L) | 61.33±8.62 | 59.67±12.58 | 60.00±7.07 | 61.67±13.01 | 89.00±5.00 | 73.00±10.00 | — |
| Lactate dehydrogenase (U/L) | 980.33±234.18 | ＞4000.00±0.00 | 732.33±73.81 | 661.67±295.19 | ＞3321.00±598.02 | ＞4000.00±0.00 | 1105-3993 |
| Creatine kinase (U/L) | 2585.67±35.99 | 3502.67±491.31 | 2221.67±108.99 | 1628.67±582.23 | 2589.33±166.77 | 1958.67±466.71 | 68-1070 |
| Creatinine (μmol/L) | 17.43±2.97 | 30.60±7.04 | 21.53±2.39 | <17.80±6.69 | 34.77±13.89 | ＜25.90±15.46 | 12.0-71.0 |
| Uric acid (μmol/L) | <21.47±19.84 | ＜22.80±12.47 | <10.00±0.00 | <10.34±0.60 | ＜117.40±173.97 | ＜247.84±171.98 | 101.0-321.0 |
| Urea (mmol/L) | 8.59±0.73 | 14.05±2.98 | 6.92±1.20 | 9.05±0.48 | 13.94±1.97 | 13.36±1.04 | 4.0-11.8 |
| Glucose (mmol/L) | 7.72±0.85 | 4.16±0.83 | 7.66±1.00 | 7.01±0.75 | 4.08±0.48 | 3.33±0.18 | 5.0-10.7 |
| Total cholesterol (mmol/L) | 2.38±0.39 | 2.15±0.28 | 2.79±0.19 | 2.37±0.24 | 1.79±0.51 | 1.90±0.17 | 0.93-4.04 |
| Triglyceride (mmol/L) | 1.16±0.17 | 0.63±0.11 | 1.86±0.52 | 1.50±0.15 | 0.74±0.13 | 0.68±0.09 | 0.62-1.63 |
| Total carbon dioxide (mmol/L) | 20.17±2.17 | 24.87±1.80 | 19.55±1.05 | 24.03±1.42 | 22.73±3.70 | 22.10±1.26 | — |
| Calcium (mmol/L) | 2.58±0.28 | 2.28±0.04 | 2.58±0.19 | 2.67±0.13 | 2.28±0.16 | 2.20±0.15 | 1.48-2.35 |
| Inorganic phosphorus (mmol/L) | 3.97±0.16 | 2.14±0.21 | 3.96±0.09 | 3.71±0.23 | 1.75±0.06 | 1.97±0.15 | 1.97-3.26 |

**Supplementary Table 10** Hematological analysis of C57BL/6J mice infected with MA-YEZV following antiviral drug treatment.

| **Test items** | **Mock** | | **YEZV** | | **YEZV+50 mg/kg Ribavirin** | **YEZV+100 mg/kg Ribavirin** | **YEZV+10 mg/kg Remdesivir** | **YEZV+30 mg/kg Remdesivir** | **Reference value** |
| --- | --- | --- | --- | --- | --- | --- | --- | --- | --- |
| White blood cell count (10^9/L) | | 3.51±1.07 | | 0.84±0.15 | 3.74±0.82 | 3.09±1.01 | 0.89±0.31 | 0.83±0.22 | 3.61-13.00 |
| Neutrophil count (10^9/L) | | 0.29±0.02 | | 0.45±0.18 | 0.33±0.07 | 0.20±0.06 | 0.56±0.29 | 0.49±0.17 | 0.10-2.00 |
| Lymphocyte count (10^9/L) | | 3.14±1.14 | | 0.16±0.04 | 3.36±0.77 | 2.84±0.98 | 0.16±0.06 | 0.17±0.10 | 1.27-8.44 |
| Monocyte count (10^9/L) | | 0.00±0.00 | | 0.14±0.13 | 0.01±0.01 | 0.01±0.01 | 0.10±0.12 | 0.08±0.08 | 0.00-0.29 |
| Eosinophil count (10^9/L) | | 0.04±0.05 | | 0.07±0.06 | 0.01±0.01 | 0.03±0.03 | 0.04±0.04 | 0.08±0.07 | 0.00-0.17 |
| Basophil count (10^9/L) | | 0.04±0.02 | | 0.02±0.01 | 0.02±0.01 | 0.01±0.01 | 0.02±0.01 | 0.02±0.02 | 0.00-0.20 |
| Neutrophil percentage (%) | | 8.80±3.38 | | 53.23±14.47 | 9.00±0.98 | 6.47±1.86 | 61.50±17.70 | 58.63±17.37 | 1.2-30.0 |
| Lymphocyte percentage (%) | | 88.10±6.59 | | 18.97±3.97 | 89.57±1.40 | 91.27±3.61 | 18.10±3.30 | 19.00±8.23 | 70.0-96.0 |
| Monocyte percentage (%) | | 0.13±0.12 | | 16.37±15.22 | 0.50±0.44 | 0.60±0.53 | 13.37±16.76 | 10.40±11.48 | 0.0-10.0 |
| Eosinophil percentage (%) | | 1.50±1.85 | | 9.13±9.84 | 0.43±0.32 | 1.17±1.17 | 5.17±4.80 | 9.43±5.40 | 0.0-10.0 |
| Basophil percentage (%) | | 1.50±1.30 | | 2.33±1.21 | 0.47±0.31 | 0.47±0.25 | 1.87±0.57 | 2.57±2.47 | 0.0-5.0 |
| Red blood cell count (10^12/L) | | 7.42±0.81 | | 8.43±0.27 | 7.05±0.49 | 9.01±1.69 | 8.37±0.30 | 8.38±1.03 | 6.00-12.50 |
| Hemoglobin concentration (g/L) | | 124.00±10.82 | | 134.00±18.25 | 120.00±4.00 | 109.67±13.61 | 134.33±8.08 | 123.67±13.58 | 100-190 |
| Hematocrit (%) | | 34.83±4.20 | | 40.20±1.44 | 36.40±1.64 | 45.97±6.30 | 40.50±0.66 | 38.87±4.52 | 40.0-48.0 |
| Mean corpuscular volume (fL) | | 47.97±1.20 | | 48.30±0.90 | 48.40±0.96 | 48.20±1.31 | 46.90±1.05 | 47.63±0.76 | 41.0-63.0 |
| Mean corpuscular hemoglobin (pg) | | 17.10±0.66 | | 17.33±0.85 | 16.33±0.74 | 12.90±3.20 | 15.67±1.00 | 15.33±1.00 | 13.0-19.0 |
| Mean corpuscular hemoglobin concentration (g/L) | | 355.33±12.58 | | 319.00±58.03 | 350.67±15.18 | 277.00±83.88 | 346.67±13.05 | 344.33±17.24 | 290-351 |
| Coefficient of variation of red cell distribution width (%) | | 17.83±0.60 | | 16.07±1.61 | 17.67±1.92 | 16.90±0.66 | 16.57±0.75 | 17.07±1.72 | 10.0-20.0 |
| Red cell distribution width standard deviation (fL) | | 33.60±1.32 | | 32.37±3.00 | 35.30±2.36 | 33.30±2.12 | 30.70±0.92 | 33.47±1.51 | 0.1-99.9 |
| Platelet count (10^9/L) | | 834.67±247.27 | | 430.00±114.14 | 747.33±73.43 | 825.67±64.90 | 474.00±55.57 | 441.33±51.52 | 540-154 |
| Mean platelet volume (fL) | | 9.50±0.20 | | 11.03±0.72 | 8.37±0.31 | 9.53±0.74 | 10.83±0.55 | 11.77±0.91 | 3.8-14.1 |
| Platelet distribution width (fL) | | 6.90±0.17 | | 9.87±2.37 | 6.37±0.76 | 13.67±7.82 | 8.80±1.39 | 10.03±1.55 | 0.1-30.0 |
| Platelet hematocrit (%) | | 0.80±0.24 | | 0.54±0.13 | 0.64±0.11 | 0.82±0.07 | 0.52±0.06 | 0.66±0.12 | 0.10-9.99 |

**Supplementary Table 11** The information of primers in this study.

| Primer | Position (bp) | Sequence (5'→3') | Polarity | Amplicon (bp) |
| --- | --- | --- | --- | --- |
| Virus Detection |  |  |  |  |
| F | 2215 | CTGCCCAATTCAAGAATC | + | 112 |
| R | 2327 | CCTCAATGTCTAGGTCTC | - |  |
| Probe | 2244 | 5'-FAM-ACAACCACTCCATCCACCACA-3'-BHQ1 | + |  |
| Genome amplification |  |  |  |  |
| L1-F | 1 | ATGGCAGGCAAGCTTTCCAAG | + | 969 |
| L1-R1 | 1166 | TGTTTAAGATCCTGCTAACCT | - |  |
| L1-R2 | 970 | AACGAATGTTCTCAGGACGTA | - |  |
| L2-F | 800 | TCAATCCCGGTGAAGCTCT | + | 845 |
| L2-R1 | 1879 | CAAACAGAACCTCGCACTC | - |  |
| L2-R2 | 1645 | GTTTGTACCAGAATACGCTCT | - |  |
| L3-F | 1533 | AAGACAAGAGCTCCCACGTA | + | 894 |
| L3-R1 | 2537 | AATCTGACAGCCTTGTACTCG | - |  |
| L3-R2 | 2427 | CCTTCCCATCTACATCGGTC | - |  |
| L4-F | 2040 | AGTGCAAACCTACTATCCTG | + | 1013 |
| L4-R1 | 3154 | TAGCGACCTAGGCTTTACTGA | - |  |
| L4-R2 | 3053 | TGGCTATACCTTTGATTGCAT | - |  |
| L5-F | 2471 | AAGAAGCCATCAAACTGTTGT | + | 960 |
| L5-R1 | 3514 | AGGCTTTCTCTGAATATCCTGT | - |  |
| L5-R2 | 3431 | TCTTAGCTGCTCTCTTTTCGG | - |  |
| L6-F | 3417 | TCCCACCAAGGTTTCCGAA | + | 818 |
| L6-R1 | 4438 | CCAGCCCAAACACAACTGA | - |  |
| L6-R2 | 4235 | AGATGCGTGTTCCTTAGTCA | - |  |
| L7-F | 4129 | ACATTCCTTATGAATCGTCGT | + | 948 |
| L7-R1 | 5285 | TTTCTAGCAAGCTCCGTGA | - |  |
| L7-R2 | 5077 | GGTTTTATCTCTGACTTTAGCCT | - |  |
| L8-F | 4953 | TAGGCATGAACTTCCAACTGA | + | 1026 |
| L8-R1 | 6103 | ATTCTTGTGCCTTAAGAGC | - |  |
| L8-R2 | 5979 | ATAGCAGCAAGAGAACTTTCG | - |  |
| L9-F | 5673 | GCTGCTCACAAATGATGGTA | + | 1035 |
| L9-R1 | 6802 | TCCTGCACTAGTAAGTCTCG | - |  |
| L9-R2 | 6708 | CTAACTGCTGAAGTACTGCC | - |  |
| L10-F | 6683 | TTAAACTCGTAGGCTGTTCCG | + | 435 |
| L10-R1 | 7544 | TCTTACTTATTGCTCCGCTGA | - |  |
| L10-R2 | 7118 | GTACATGTTGCACCAATCCG | - |  |
| L11-F | 6826 | GGCACTAAAGTAGTCCATGCAA | + | 531 |
| L11-R1 | 7680 | GTAAAACTCCAGCATTGTGTCC | - |  |
| L11-R2 | 7357 | GTTGTAACATGTCATCGCCAT | - |  |
| L12-F | 7306 | GACCAGCCCATGATACAGT | + | 1083 |
| L12-R1 | 8441 | ATCTAACAAGGCCAAATGTCC | - |  |
| L12-R2 | 8389 | ACAAACTCAAAGTTAGACCGGA | - |  |
| L13-F | 8839 | AGGATTATGACGGATTCCCT | + | 426 |
| L13-R1 | 9357 | GGCCTCCGTAATCAACCCTG | - |  |
| L13-R2 | 9265 | TTGACCTGCTCAGATGCATT | - |  |
| L14-F | 9157 | ATGCAGTTTTCAGAGCCGAA | + | 1115 |
| L14-R1 | 10311 | GTGTAAAGCCCAACATCCTT | - |  |
| L14-R2 | 10272 | GACTCCCCTGATGTACCCA | - |  |
| L15-F | 10230 | CCTGCTTCCAACCTATCCAA | + | 921 |
| L15-R1 | 11285 | GAATCATATCATTGCCGCTGT | - |  |
| L15-R2 | 11151 | AGACCTCTGTCATCTAGGCTG | - |  |
| L16-F1 | 11190 | CACTTTGTACCAGCTACAGG | + | 534 |
| L16-F2 | 111264 | TAGTGACCAGCAAGACCTT | + |  |
| L16-R | 11798 | TAGAAACTCAGATCAAACAG | - |  |
| M1-F | 1 | ATGTTAATCATGCAGTGTAG | + | 787 |
| M1-R1 | 859 | ATTGTTCACCGTGTTGACCTG | - |  |
| M1-R2 | 788 | TGCCTTACAGCCTCTTGCAT | - |  |
| M2-F | 641 | CCGGATGTGAGAAATTACACAA | + | 1001 |
| M2-R1 | 1692 | GTCAAGCGGTTAAAGATCCAC | - |  |
| M2-R2 | 1642 | CTACCAATGCTGTAGCCGAA | - |  |
| M3-F | 1640 | TTTTCGGCTACAGCATTGGTA | + | 727 |
| M3-R1 | 2417 | CACTTCAGCCCAGACCGAG | - |  |
| M3-R2 | 2367 | CAGCCACCACAACCTTGCTA | - |  |
| M4-F | 2322 | GGGCAACCACATTAGCCTC | + | 1059 |
| M4-R1 | 3447 | ACAAGTTATACCTGTGGCGCAA | - |  |
| M4-R2 | 3381 | GCAGCACTGTCTTTGACTCT | - |  |
| M5-F1 | 3136 | ACTGTGCTTCAATACTACTGCAA | + | 829 |
| M5-R2 | 3210 | AGGTGCGTTTGAAAACCTCT | + |  |
| M5-R | 4039 | ATCGTTCCTTCTCAGCAGTTC | - |  |
| S1-F | 1 | ATGGCACGTCTGATTGAGGC | + | 766 |
| S1-R1 | 798 | TCCATCTTCTGCTGGTTGTTG | - |  |
| S1-R2 | 767 | TGCCTTTTCCACCTTTGACC | - |  |
| S2-F | 478 | ATTGCCCCTAAGAAATACGTGA | + | 526 |
| S2-R1 | 1119 | TTGAAGTCTCCTGTTGCACAC | - |  |
| S2-R2 | 1004 | CATCTCATTCAGGGCACCAA | - |  |
| S3-F1 | 789 | AGAACTTCTCAACAACCAGC | + | 557 |
| S3-F2 | 935 | GTTCAGCCCTTGACACTGC | + |  |
| S3-R | 1492 | TTAAGGTTTTCCTCCAGT | - |  |
| Cytokine detection |  |  |  |  |
| IFNγ-F |  | TCAAGTGGCATAGATGTGGAAGAA | + |  |
| IFNγ-R |  | TGGCTCTGCAGGATTTTCATG | - |  |
| TNFɑ-F |  | CTGTGAAGGGAATGGGTGTT | + |  |
| TNFɑ-R |  | GGTCACTGTCCCAGCATCTT | - |  |
| IL1β-F |  | CGCAGCAGCACATCAACAAGAGC | + |  |
| IL1β-R |  | TGTCCTCATCCTGGAAGGTCCACG | - |  |
| IL6-F |  | CTGCAAGAGACTTCCATCCAG | + |  |
| IL6-R |  | AGTGGTATAGACAGGTCTGTTGG | - |  |
| IL10-F |  | CTTACTGACTGGCATGAGGATCA | + |  |
| IL10-R |  | GCAGCTCTAGGAGCATGTGG | - |  |
| IL12-F |  | CAATCACGCTACCTCCTCTTTT | + |  |
| IL12-R |  | CAGCAGTGCAGGAATAATGTTTC | - |  |
| IL18-F |  | GCCATGTCAGAAGACTCTTGCGTC | + |  |
| IL18-R |  | GTACAGTGAAGTCGGCCAAAGTTGTC | - |  |
| NLRP1-F |  | CTCACATCCACATACTGCTCAC | + |  |
| NLRP1-R |  | CAACATCTTCACACCACCATCA | - |  |
| NLRP3-F |  | CCAGACCTCCAAGACCACTAC | + |  |
| NLRP3-R |  | TACATAGCAGCGAAGAACTCCT | - |  |
| NLRP4-F |  | CAAGGCTCTGCTGCTGAAG | + |  |
| NLRP4-R |  | CGTGGTGGTGGTGACAATG | - |  |
| AIM2-F |  | TGGCAGATAGGACAGAGTTAGC | + |  |
| AIM2-R |  | CAGAGGCAGCAGAGCAGTT | - |  |
| CCL2-F |  | AAGAGATCAGGGAGTTTGCT | + |  |
| CCL2-R |  | CTGCCTCCATCAACCACTTT | - |  |
| CCL3-F |  | TTCTCTGTACCATGACACTCTGC | + |  |
| CCL3-R |  | CGTGGAATCTTCCGGCTGTAG | - |  |
| CCL4-F |  | TTCCTGCTGTTTCTCTTACACCT | + |  |
| CCL4-R |  | CTGTCTGCCTCTTTTGGTCAG | - |  |
| CCL5-F |  | GCTGCTTTGCCTACCTCTCC | + |  |
| CCL5-R |  | TCGAGTGACAAACACGACTGC | - |  |
| CCL11-F |  | GAATCACCAACAACAGATGCAC | + |  |
| CCL11-R |  | ATCCTGGACCCACTTCTTCTT | - |  |
| CXCL1-F |  | CTGGGATTCACCTCAAGAACATC | + |  |
| CXCL1-R |  | CAGGGTCAAGGCAAGCCTC | - |  |
| CXCL2-F |  | TGAACAAAGGCAAGGCTAACTG | + |  |
| CXCL2-R |  | AAGTGAACTCTCAGACAGCGAGG | - |  |
| CXCL5-F |  | TGGGCAGTGACAAAAAGAAAGC | + |  |
| CXCL5-R |  | AAATCCGTGGGTGGAGAGAATC | - |  |
| CXCL9-F |  | TGCTACACTGAAGAACGGAGAT | + |  |
| CXCL9-R |  | TCCTTGAACGACGACGACTT | - |  |
| CXCL10-F |  | GCTGCAACTGCATCCATATCG | + |  |
| CXCL10-R |  | CATCGTGGCAATGATCTCAACA | - |  |
| CASP1-F |  | ATGAAGTTGCTGCTGGAGGAT | + |  |
| CASP1-R |  | ACTGTCAGAAGTCTTGTGCTCT | - |  |
| CASP3-F |  | GCTGACTTCCTGTATGCTTACT | + |  |
| CASP3-R |  | ATTCCGTTGCCACCTTCCT | - |  |
| CASP6-F |  | AACGCAGACAGAGACAACCT | + |  |
| CASP6-R |  | TCGGCATCTATGTGGCTTGA | - |  |
| CASP7-F |  | TTGCTTACTCCACGGTTCCA | + |  |
| CASP7-R |  | GGTCCTTGCCATGCTCATTC | - |  |
| CASP8-F |  | GCCTCCATCTATGACCTGACA | + |  |
| CASP8-R |  | GTTCTTCACCGTAGCCATTCC | - |  |
| CASP12-F |  | TCCTCAGACAGCACATTCCT | + |  |
| CASP12-R |  | TTCTCAGACTCCGACAGTTAGA | - |  |
| GSDMC-F |  | CCTGATGGTGCTGAGTGATAC | + |  |
| GSDMC-R |  | GCTGAGGCTGGAGTGTGAA | - |  |
| GSDMC-F |  | ATACTGCGTGTGACTCAGAAGA | + |  |
| GSDMC-R |  | CTCGGAATGCCAGGATGCT | - |  |
| GSDME-F |  | GATGTGGCTGGTGGTCTCT | + |  |
| GSDME-R |  | GCTCTCCTTGTATCCTGTATCC | - |  |
| ACTB-F |  | GAGACCTTCAACACCCCAGC | + |  |
| ACTB-R |  | ATGTCACGCACGATTTCCC | - |  |
